# Walrus Population-specific Marine Reservoir Offsets (ΔR) for Calibration of Radiocarbon Dates: Implications for Arctic Chronologies and Medieval Trade

**DOI:** 10.64898/2026.01.21.698949

**Authors:** Bente Philippsen, Katrien Dierickx, Mohsen Falahati-Anbaran, Erin H. Kunisch, Thomas C.A. Royle, Richard Sabin, Eirik Sollid, James H. Barrett

## Abstract

Walruses have long played a vital role in Arctic subsistence and commercial economies, yet accurate radiocarbon dating of walrus-derived archaeological materials is complicated by regional variation in marine reservoir effects. This study presents new ΔR values derived from 31 new and 15 legacy radiocarbon dates on known-age walrus specimens spanning multiple populations of *Odobenus rosmarus rosmarus* and *O. r. divergens*. The results reveal substantial inter-population variability, with ΔR values ranging from −140 (Franz Josef Land) to +295 (Pacific), underscoring the importance of population-specific calibration. A pooled ΔR of +17±12 was calculated for western Greenland and the Canadian Arctic—regions central to the medieval Norse ivory trade—and applied to 12 walrus rostra and ivory artefacts excavated in Trondheim, Norway. The resulting calibrated dates are from the 11th to early 14th centuries CE, confirming that seemingly anomalous post-15th-century finds are residual rather than evidence of continued trade. An alternative ΔR estimate of −110±35, derived solely from archaeological context dates, suggests potential time lags between harvest and deposition. These findings demonstrate the value of known-age walrus ΔR data for refining chronologies of Arctic exploitation and long-distance trade, while highlighting the need for provenance studies in archaeological dating and both the pertinence and limitations of mollusk-based reservoir corrections.

## Introduction

Walruses have been an important resource for northern communities for centuries. They provided meat, blubber, hide and/or ivory (Frei et al. 2015; Mason and Rasic 2019; Keighley et al. 2021). Today, they remain a valued food item and focus of small-scale subsistence hunts among Chukchi, Inuit, and Yupik communities in Greenland, Arctic Canada, Alaska and Russia (Born et al. 2017; Desjardins and Gotfredsen 2021; Metcalf et al. 2025; Zdor et al. 2010). Historically, walruses were commercially exploited on a large scale by European hunters, whalers and settlers. This led to severe population declines and in some instances extirpations (e.g. Kovacs et al. 2014; McLeod et al. 2014; Keighley et al. 2019; Barrett et al. 2020; McCaffrey 2020; Gjertz 2021).

Research on the cultural role of walrus hunting, human and environmental impacts on walrus populations and their ecosystem often relies on absolute chronologies derived from radiocarbon dating. These dates are calibrated using a calibration curve that accounts for the marine reservoir effect (e.g. Dyke et al. 2019; Dury et al. 2021). However, the standard marine calibration curve, Marine20, represents a global modelled ocean and corrects only for the average reservoir age, R. It is not intended for calibration in polar regions (Heaton et al. 2020). Reservoir ages vary locally due to factors such as upwelling of older bottom water, inflow of carbonate-rich river water, and seasonal ice cover, which limits CO_2_ exchange with the atmosphere. This variability is especially pronounced in the Arctic Ocean, where a stable surface layer and permanent ice cover further restrict exchange with atmospheric CO_2_(Heaton et al. 2020, Mangerud and Gulliksen 1975). Conversely, areas with shallow water and better atmospheric exchange may have lower reservoir ages. The local deviation from the global modelled reservoir age is denoted ΔR and is essential for accurate calibration of samples of unknown age.

To estimate ΔR, known-age samples from specific localities are dated so that their local offset can be calculated. Samples collected before the 1950s are preferred to avoid effects from the ^14^C excess produced by atomic bomb tests. In this paper, we discuss and calculate ΔR values respective to the Marine20 curve, despite its limited suitability for polar regions, to ensure compatibility with recent publications and databases, such as the 14CHRONO Marine20 Reservoir database (calib.org/marine, Reimer and Reimer 2001). When we refer to ΔR values respective to other marine calibration curves, we will specify this, e.g. as ΔR_Marine13_. Relatively extensive ΔR datasets are available from mollusk shells, as these are easy to collect, curate, and sample after decades of storage (e.g. Reimer and Reimer 2001; Pieńkowski et al. 2023; Pearce et al. 2023). However, mollusk shells may not be representative of walrus diet, even when mollusks dominate the diet. The flesh of the mollusks can have ΔR values that differ from those of their shells (Erlenkeuser et al. 1975: 288-290; Fernandes et al. 2012). Walrus dentine and bone collagen are formed from dietary protein rather than shell carbonate (Dyke et al. 2019, Jim et al. 2004). For example, the δ^13^C value of mollusk shell was shown to change throughout ontogeny in one species, indicating shifts in carbon source for shell formation (Lorrain et al. 2004). The carbon in the flesh derives from the mollusk’s diet, while the majority of the carbon in the shell carbonate derives from water dissolved inorganic carbon, or DIC (McConnaughey and Gillikin 2008). In many cases, the age difference should not be significant, as the diet of e.g. filter feeders may be dominated by phytoplankton, whose DIC source is the same as that for the mollusk shell (Gillikin et al. 2006). In this paper, we radiocarbon date bones of walruses with known years of death to avoid such possible ambiguities. These walruses are from different populations or regional groupings previously identified on the basis of genetic and other evidence (e.g. Keighley et al. 2022; Ruiz-Puerta 2023; Dierickx et al. forthcoming), to calculate their ΔR. The focus is on populations of the Atlantic subspecies, *Odobenus rosmarus rosmarus*, for which 43 known-age specimens pre-dating 1950 were available in natural history museums for radiocarbon dating (comprising 29 new and 15 previously studied samples, with one overlap). The resulting ΔR values are divided by population where practicable. A pooled value for Greenland (excluding East Greenland) and the Canadian Arctic is also calculated for use with walrus artefacts traded to Europe for which a precise location of harvest cannot be inferred. For comparison with future work, a more tentative ΔR is also provided for the Pacific subspecies, *Odobenus rosmarus divergens*, based on two known-age specimens.

As a case study, we employ the ΔR results to interpret the chronology of medieval artefacts of walrus rostrum (skull) bone and tusk ivory recovered from archaeological contexts in Trondheim, Norway, and its hinterland. Trondheim was one of several major hubs for the trade and carving of walrus tusks, which were traded as pairs attached to the rostrum bone (Barrett 2021; Barrett and Grav 2025). Ancient DNA (aDNA) and stable isotope evidence shows that the material originated in western Greenland and/or Arctic Canada, indicating that it was acquired from or via the Norse colony of Greenland (Barrett et al. 2020; Ruiz-Puerta et al. 2024). Two questions are asked: (a) whether the pooled ΔR for Greenland and the Canadian Arctic based on known age material from natural history collections matches a ΔR estimated based on stratified archaeological finds from Trondheim and (b) whether walrus artefacts from the town that seem to post-date the 15th-century Norse abandonment of Greenland based on archaeological evidence are residual finds from earlier centuries, or if existing interpretations of the chronology of the Greenlandic trade in walrus ivory should instead be reassessed?

## Materials and Methods

Samples of bone from 31 known-age walrus specimens pre-dating 1950 were collected from the American Museum of Natural History (New York, NY, USA; *n*=10), the Canadian Museum of Nature (Ottawa, ON, CA, and Gatineau, QC, CA; *n*=4), the Natural History Museum (London, UK; *n*=7) and the Zoological Museum, Natural History Museum of Denmark (Copenhagen, DK; *n*=10). Locations and dates of harvest were available from the catalogue records of the relevant museums, and were sometimes also preserved on original labelling (Figure 1A, Figure 2). Placenames indicating catch locations represented past usage, but modern locations could be inferred with confidence (Table 1). The finds derive from Arctic exploration between 1852/53 and 1929, including expeditions by (for example) Knud Rasmussen/Peter Freuchen (5th Thule expedition to Nunavut), F.G. Jackson (Franz Josef Land), Alwin Pedersen (Scoresby Sund/Kangertittivaq) and Robert Peary (north-west Greenland).

**Table 1.** New radiocarbon dates on walrus specimens with historical dates of death. Full specimen details and museum catalogue numbers are available in Supplemental material Table S1. European place-names are retained if needed to avoid ambiguity.

| Lab number (TRa-) | Population | Origin as historically catalogued | Origin | Date of death CE | <sup>14</sup>C Age BP | <sup>14</sup>C error (1σ) |
| --- | --- | --- | --- | --- | --- | --- |
| 22945 | Baffin Bay | Wellington Channel, Nunavut | Wellington Channel | 1852-1853 | 649 | 12 |
| 22943 | Baffin Bay | Barrow Strait, Nunavut | Barrow Strait | 1853-1854 | 725 | 15 |
| 22944 | Baffin Bay | Barrow Strait, Nunavut | Barrow Strait | 1854 | 737 | 13 |
| 25066 | Baffin Bay | Murchison Sound, Greenland | Murchison Sund | 1895 | 582 | 13 |
| 25068 | Baffin Bay | Manson Island, Greenland | Qeqertaarsuit (Manson Øer) | 1896 | 603 | 14 |
| 25065 | Baffin Bay | Manson Island, Greenland | Qeqertaarsuit (Manson Øer) | 1896 | 599 | 15 |
| 25069 | Baffin Bay | Manson Island, Greenland | Qeqertaarsuit (Manson Øer) | 1896 | 582 | 13 |
| 25064 | Baffin Bay | Murchison Sound, Greenland | Murchison Sund | 1896 | 535 | 13 |
| 25063 | Baffin Bay | Murchison Sound, Greenland | Murchison Sund | 1896 | 569 | 21 |
| 25062 | Baffin Bay | Northumberland Island, Greenland | Kiatak | 1926 | 565 | 18 |
| 25061 | Baffin Bay | Northumberland Island, Greenland | Kiatak | 1926 | 563 | 13 |
| 25060 | Baffin Bay | Northumberland Island, Greenland | Kiatak | 1926 | 553 | 16 |
| 25067 | Western Greenland | Holsteinsborg, Greenland | Sisimiut | 1895 | 611 | 20 |
| 25591 | Western Greenland | Holsteinsborg, Greenland | Sisimiut | 1898 | 604 | 13 |
| 25595 | Foxe Basin | Frozen Strait, Nunavut | Frozen Strait | 1922 | 646 | 18 |
| 25596 | Foxe Basin | Frozen Strait, Nunavut | Frozen Strait | 1922 | 687 | 16 |
| 24149 | Hudson Strait/ Bay | Nottingham Island, Nunavut | Tujjaat | 1884–1885 | 671 | 20 |
| 24150 | Hudson Strait/ Bay | Prince of Wales Sound, Hudson Strait, Nunavut | Hudson Strait | 1884–1885 | 639 | 13 |
| 24148 | Hudson Strait/ Bay | Ashe Inlet, Hudson Strait, Nunavut | Ashe Inlet | 1886 | 609 | 12 |
| 24151 | Hudson Strait/ Bay | Fort Churchill, Manitoba | Churchill | 1910 | 615 | 12 |
| 25589 | Eastern Greenland | North-east Greenland | Tunu | 1906–1909 | 631 | 21 |
| 25593 | Eastern Greenland | Scoresby Sund, Greenland | Kangertittivaq | 1924 | 569 | 13 |
| 25587 | Eastern Greenland | Scoresby Sund, Greenland | Kangertittivaq | 1927 | 605 | 23 |
| 25588 | Eastern Greenland | Scoresby Sund, Greenland | Kangertittivaq | 1927–1929 | 558 | 16 |
| 25585 | Eastern Greenland | Scoresby Sund, Greenland | Kangertittivaq | 1929 | 561 | 16 |
| 25586 | Eastern Greenland | Scoresby Sund, Greenland | Kangertittivaq | 1929 | 567 | 15 |
| 25590 | Iceland | Þorlákshöfn, Iceland | Þorlákshöfn | 1900 | 576 | 14 |
| 22952 | Franz Joseph Land | Cape Flora, Franz Josef Land | Cape Flora | 1894–1897 | 472 | 15 |
| 22947 | Franz Joseph Land | Franz Josef Land | Franz Josef Land | 1894–1897 | 494 | 15 |
| 22951 | Pacific | Arctic Ocean (75° 10' N Lat; 168° 24' W Long) | Arctic Ocean | 1865 | 843 | 13 |
| 22950 | Pacific | Kamchatka | Kamchatka | 1896–1897 | 1026 | 16 |

**Figure 1.**
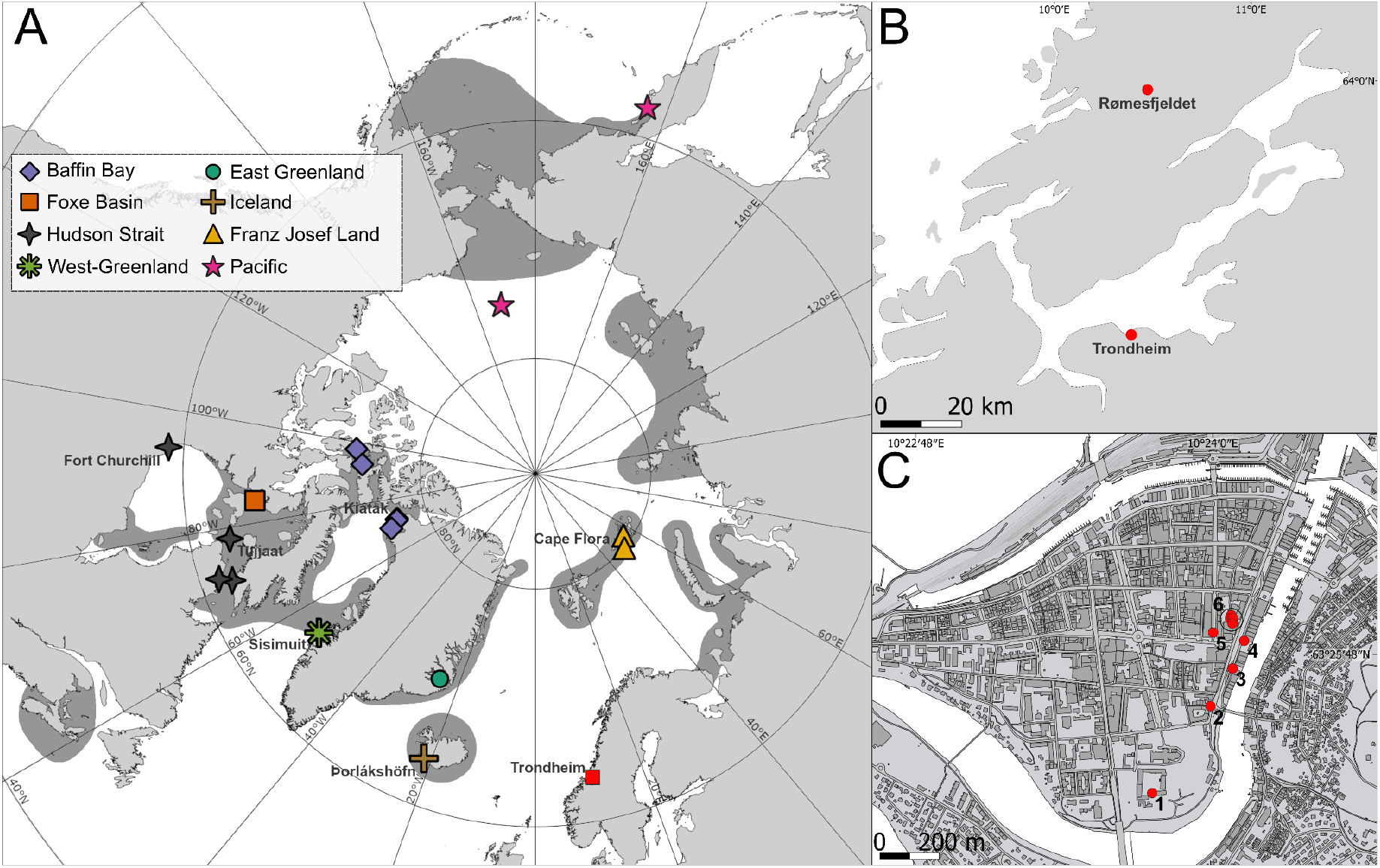
Geographical locations of: A) newly radiocarbon dated walrus specimens from natural history collections with known date of death, plotted in relation to the modern circumpolar distribution of the species (dark grey); B) Trondheim and Rømesfjeldet, the latter being the find spot of a walrus tusk radiocarbon dated in this study; C) excavations within Trondheim yielding finds of walrus rostra and/or walrus ivory that are radiocarbon dated in this study (1 Erkebispegården, 2 Kjøpmannsgata, 3 Kjøpmannsgata Waterfront, 4 Kjøpmannsgata 25-27, 5 Søndre gate, 6 Folkebibliotekstomten). Modern walrus distribution based on Andersen et al. (2014), McLeod et al. (2014), Keighley et al. (2019), Beatty et al. (2020), Keighley et al. (2022) and Mills et al. (2024).

**Figure 2.**
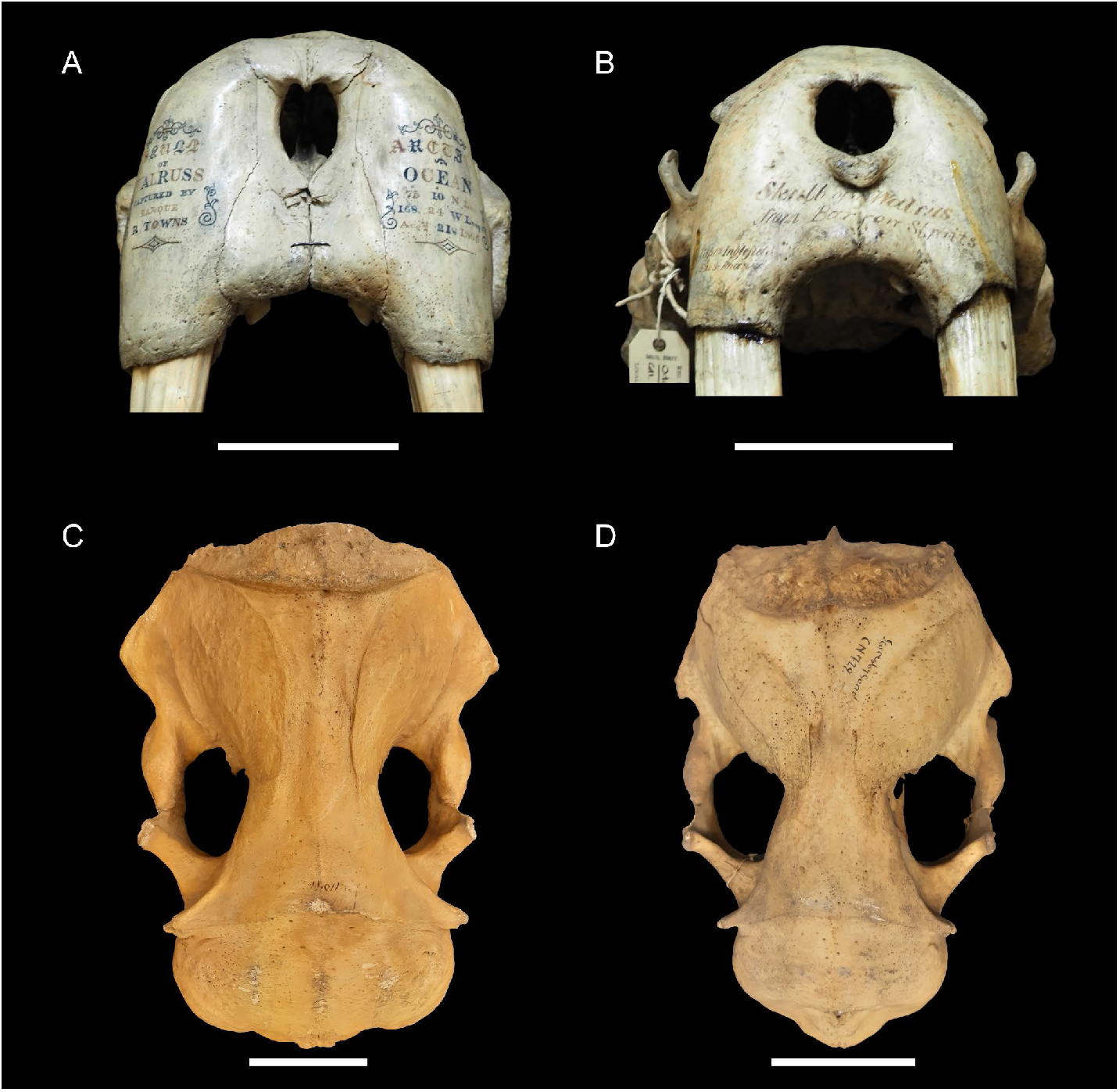
Examples of walrus specimens from natural history collections having known dates of death: TRa numbers 22951 (A, NHMUK GERM 331.j) and 22943 (B, NHMUK ZD.1855.11.26.38) from the collections of the Natural History Museum, London, 25065 (C, AMNH MO-11049) from the collections of the American Museum of Natural History, and 25588 (D, NHMD-M11-CN729) from the collections of the Natural History Museum of Denmark (photographs by James Barrett (A, B) and Katrien Dierickx (C, D)). Scale bars indicate 10 cm.

For the Trondheim case study, six samples of walrus skull (rostrum) bone and six samples of walrus ivory (one crucifix corpus, one gaming piece, three offcuts of raw material and one complete tusk) were obtained from archaeological collections curated at the NTNU University Museum (Trondheim, NO) (Table 2, Figure 3). The rostra represent the original ‘packaging’ in which pairs of tusks were traded from Greenland in the Middle Ages (Barrett et al. 2020). Eleven of the Trondheim area finds were from within the medieval town itself, with the last sample being from one of two tusks (from the same animal) bearing owner’s markings that were found at Rømesfjeldet (Rømmen, Åfjorden) north of the town (Figure 1B-1C).

**Table 2.**
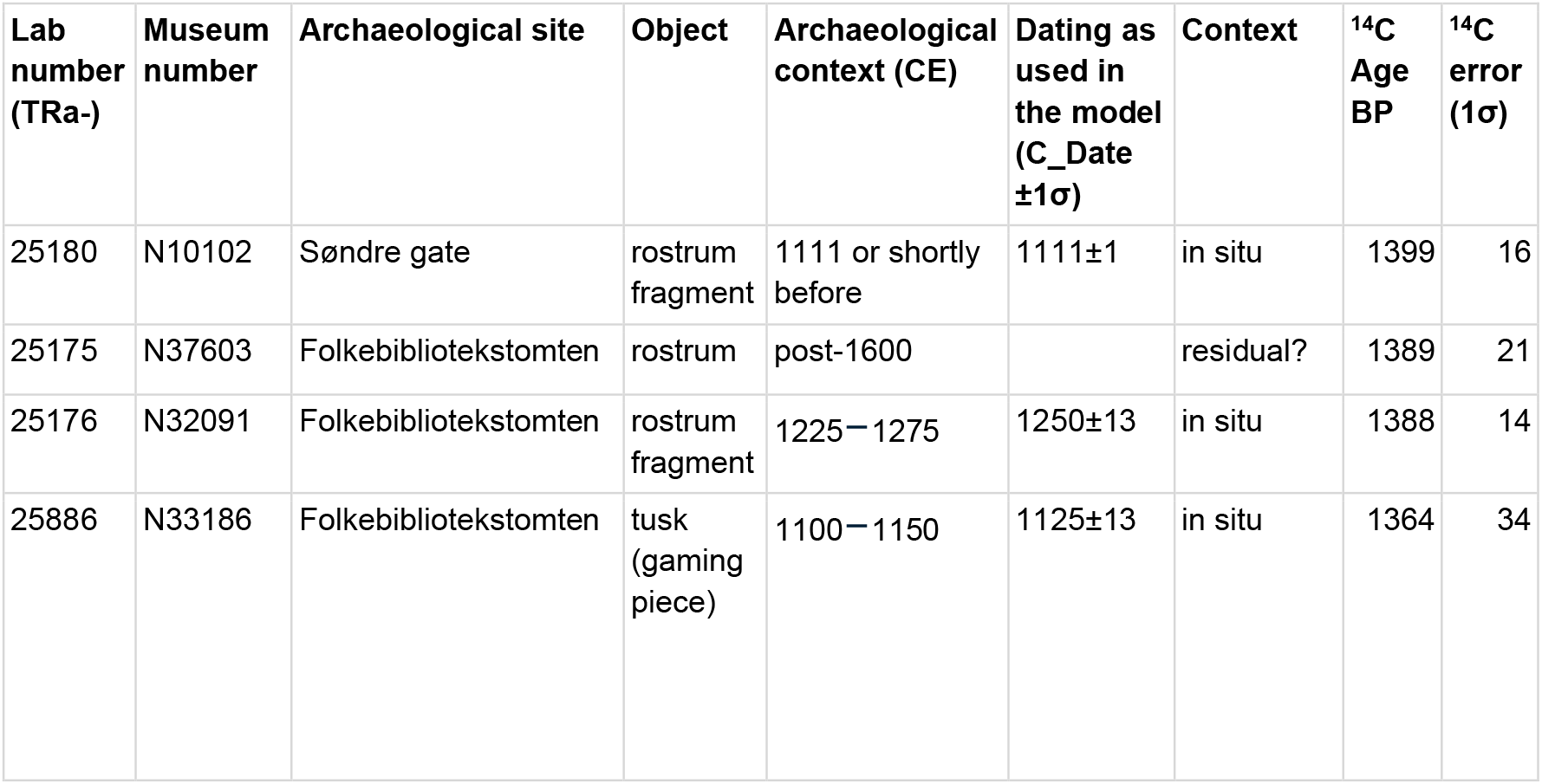
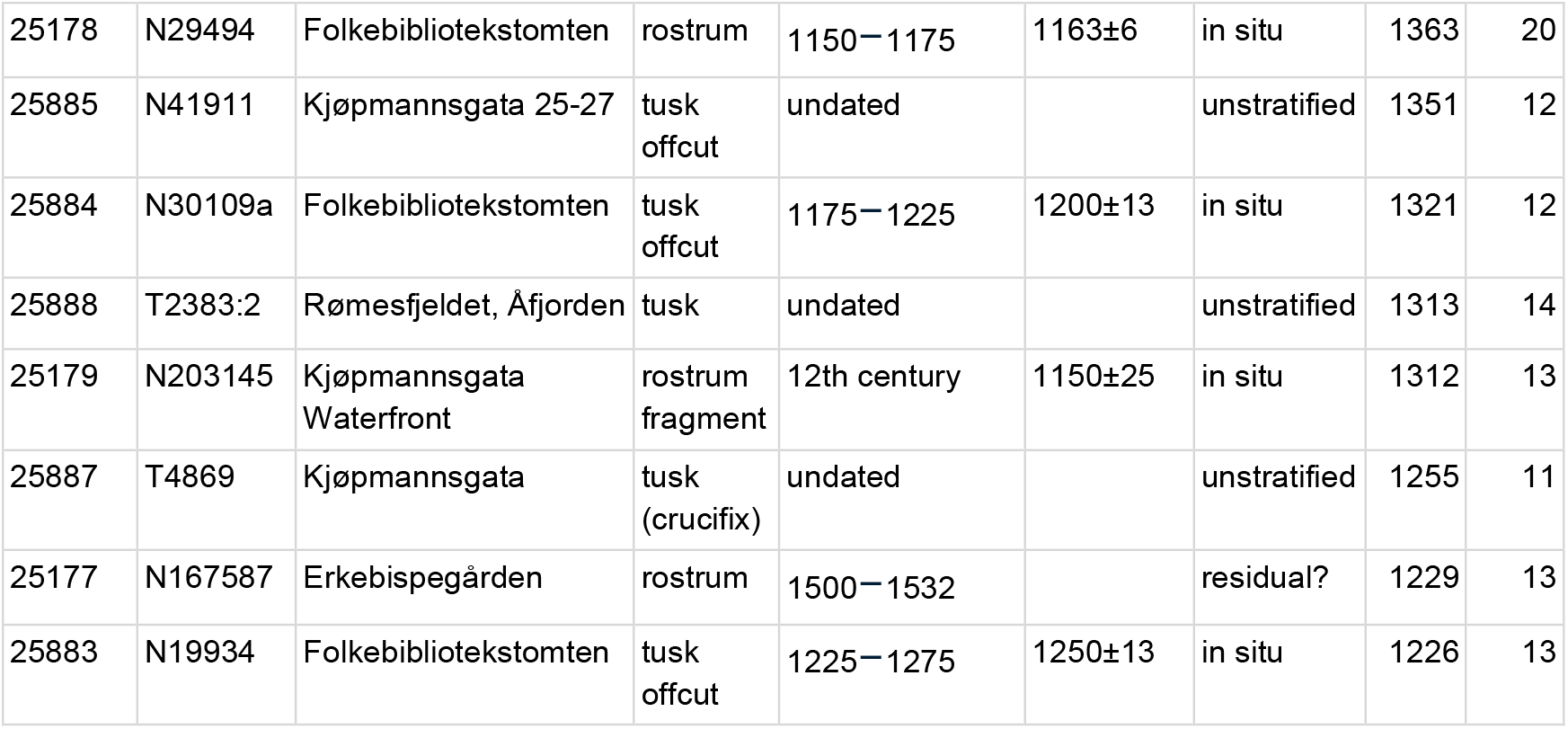
Radiocarbon dated walrus specimens from excavations in Trondheim and its hinterland. Full specimen details are available in Supplemental material Table S2. The dates are here calibrated (95.4% range) with the ΔR of +17±12 calculated for historical specimens from Greenland (excluding Eastern Greenland) and the Canadian Arctic.

| Lab number (TRa-) | Museum number | Archaeological site | Object | Archaeological context (CE) | Dating as used in the model (C_Date $\pm 1\sigma$) | Context | <sup>14</sup>C Age BP | <sup>14</sup>C error (1σ) |
| --- | --- | --- | --- | --- | --- | --- | --- | --- |
| 25180 | N10102 | Søndre gate | rostrum fragment | 1111 or shortly before | 1111$\pm 1$ | in situ | 1399 | 16 |
| 25175 | N37603 | Folkebibliotekstomten | rostrum | post-1600 |  | residual? | 1389 | 21 |
| 25176 | N32091 | Folkebibliotekstomten | rostrum fragment | 1225–1275 | 1250$\pm 13$ | in situ | 1388 | 14 |
| 25886 | N33186 | Folkebibliotekstomten | tusk (gaming piece) | 1100–1150 | 1125$\pm 13$ | in situ | 1364 | 34 |
| 25178 | N29494 | Folkebibliotekstomten | rostrum | 1150–1175 | 1163±6 | in situ | 1363 | 20 |
| 25885 | N41911 | Kjøpmannsgata 25-27 | tusk<br>offcut | undated |  | unstratified | 1351 | 12 |
| 25884 | N30109a | Folkebibliotekstomten | tusk<br>offcut | 1175–1225 | 1200±13 | in situ | 1321 | 12 |
| 25888 | T2383:2 | Rømesfjeldet, Åfjorden | tusk | undated |  | unstratified | 1313 | 14 |
| 25179 | N203145 | Kjøpmannsgata<br>Waterfront | rostrum<br>fragment | 12th century | 1150±25 | in situ | 1312 | 13 |
| 25887 | T4869 | Kjøpmannsgata | tusk<br>(crucifix) | undated |  | unstratified | 1255 | 11 |
| 25177 | N167587 | Erkebispegården | rostrum | 1500–1532 |  | residual? | 1229 | 13 |
| 25883 | N19934 | Folkebibliotekstomten | tusk<br>offcut | 1225–1275 | 1250±13 | in situ | 1226 | 13 |

**Table 3.** ΔR values per population (95.4% ranges) based on differing OxCal models and the exclusion or inclusion of legacy radiocarbon dates; n = number of dated specimens. BAF Baffin Bay, WGL West Greenland, FOX Foxe Basin, HUD Hudson Strait and Hudson Bay, EGL East Greenland, ICE Iceland, FJL Franz Josef Land, PAC Pacific (see Figure 1). DoD Date of death, eDoB estimated date of birth.

| Population | OxCal constraints | $\Delta R$ range from | to | $\Delta R$ mean | $\Delta R$ 1stdev | Dates included |
| --- | --- | --- | --- | --- | --- | --- |
| BAF (n=12) | DoD | -98 | +2 | -47 | 24 | This study |
| BAF (n=12) | eDoB & DoD | -66 | +14 | -26 | 19 | This study |
| BAF (n=13) | eDoB & DoD | -51 | +27 | -12 | 19 | This study; Dyke et al. 2019 |
| WGL (n=2) | DoD | -199 | +81 | -54 | 72 | This study |
| WGL (n=2) | eDoB & DoD | -113 | +78 | -17 | 47 | This study |
| FOX (n=2) | DoD | -215 | +169 | +13 | 95 | This study |
| FOX (n=2) | eDoB & DoD | -31 | +156 | +62 | 46 | This study |
| FOX (n=5) | eDoB & DoD | -10 | +109 | +49 | 29 | This study; Dyke et al. 2019 |
| HUD (n=4) | DoD | -110 | +61 | -23 | 43 | This study |
| HUD (n=4) | eDoB & DoD | -63 | +70 | +4 | 33 | This study |
| HUD (n=14) | eDoB & DoD | 0 | +71 | +35 | 18 | This study; Dyke et al. 2019 |
| Pooled $\Delta R$ for Greenland and Canadian Arctic (BAF, WGL, FOX, HUD; n=20) | DoD | -72 | +5 | -33 | 19 | This study |
| Pooled $\Delta R$ for Greenland and Canadian Arctic (BAF, WGL, FOX, HUD; n=20) | eDoB & DoD | -40 | +20 | -10 | 15 | This study |
| Pooled $\Delta R$ for Greenland and Canadian Arctic (BAF, WGL, FOX, HUD; n=34) | eDoB & DoD | -9 | +42 | +17 | 12 | This study; Dyke et al. 2019 |
| EGL (n=6) | DoD | -119 | +20 | -48 | 33 | This study |
| EGL (n=6) | eDoB & DoD | -82 | +29 | -26 | 27 | This study |
| ICE (n=1) | DoD | -408 | +85 | -115 | 121 | This study |
| ICE (n=1) | eDoB & DoD | -182 | +84 | -49 | 66 | This study |
| FJL (n=2) | DoD | -391 | -42 | -189 | 82 | This study |
| FJL (n=2) | eDoB & DoD | -235 | -45 | -140 | 47 | This study |
| PAC (n=2) | DoD | -103 | +387 | +179 | 129 | This study |
| PAC (n=2) | eDoB & DoD | +200 | +391 | +295 | 47 | This study |

**Figure 3.**
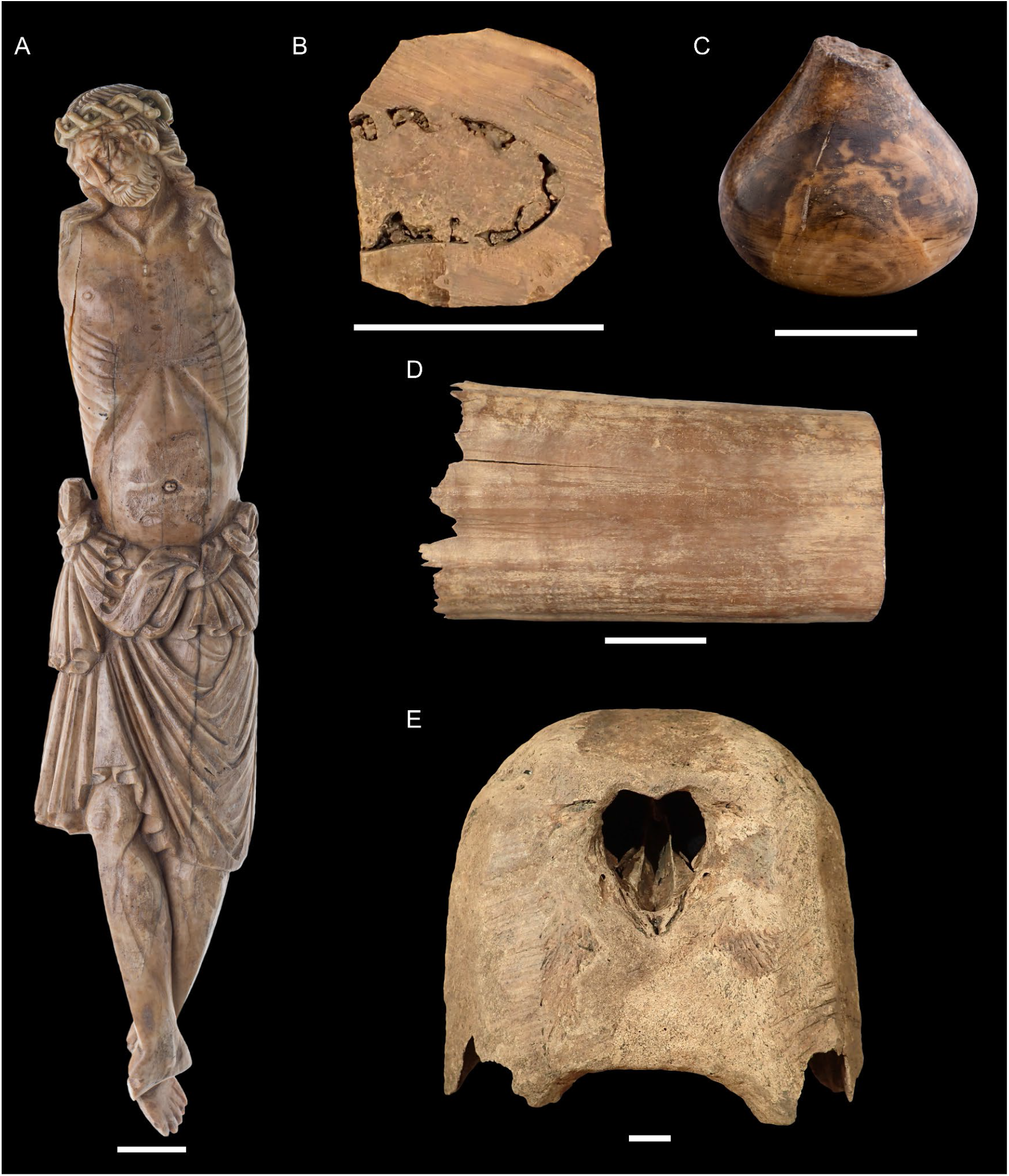
A selection of the medieval walrus ivory artefacts and rostrum bone from Trondheim radiocarbon dated in this study: museum numbers T4869 (A, crucifix), N30109a (B, tusk offcut), N33186 (C, gaming piece), N41911 (D, tusk offcut), N37603 (E, rostrum), (photographs by Åge Hojem (A, C) and J.H. Barrett (B, D, E), NTNU University Museum). Scale bars indicate 2 cm.

**Figure 4.**
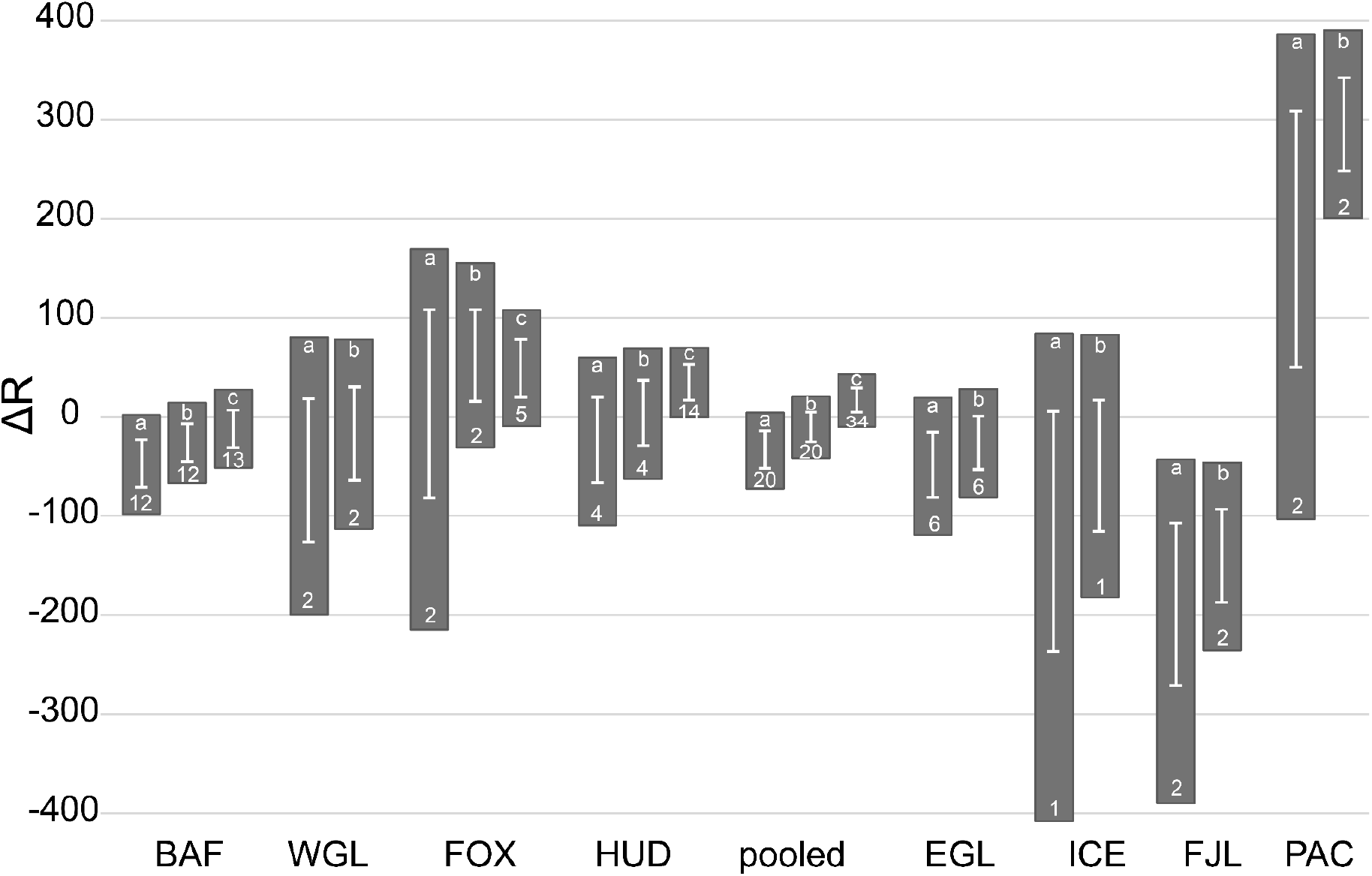
ΔR values per population (95.4% ranges) based on differing OxCal models and the exclusion or inclusion of legacy radiocarbon dates; data from Table 3. BAF Baffin Bay, WGL West Greenland, FOX Foxe Basin, HUD Hudson Strait and Hudson Bay, EGL East Greenland, ICE Iceland, FJL Franz Josef Land, PAC Pacific (see Figure 1). The error bar denotes the mean ΔR plus/minus one standard deviation; the box marks the ΔR range (from Table 3). The letters in the upper part of the boxes mark the calculation: a) only using date of death, b) additionally using estimated date of birth, c) using both date of death, estimated date of birth, and data from Dyke et al. (2019). The numbers in the lower parts of the boxes are the numbers of samples included in these averages.

The newly studied known-age natural history specimens were sampled using a Dremel rotary tool, either by cutting a small chunk from a cranium using a diamond rotary blade (n=28) or drilling powder from a mandibular tooth socket using a steel bit and/or a diamond bur (n=3). Most (n=19) of the cranial samples were cut from tympanic bullae, with the remainder (n=9) taken from other locations (mastoid, parietal, temporal, tusk socket or other maxillary bone) depending on the completeness of the specimens and the guidance of museum curators. Original samples were typically between 0.5g and 1g, as agreed with the relevant museum and/or heritage authority. The archaeological samples from the NTNU University Museum were also sampled with a Dremel rotary tool using a diamond rotary blade (to cut a small chunk) or diamond burrs and steel drill bits (to extract powder). For these artefacts, curatorial considerations sometimes necessitated smaller samples of c. 0.3–0.4g. Both the natural history and archaeological samples were used for multiple analyses (e.g. isotope analysis and ancient DNA (aDNA)) of which the radiocarbon dating reported here was one. Potentially contaminated (e.g. dusty) external surfaces were abraded or cut away prior to on-site sampling or in-lab subsampling.

Collagen was extracted from the bone or ivory samples with a modified Longin procedure (Longin 1971) as described in Seiler et al. (2019), with additional modifications to maximise yield and minimise potential contamination. Samples were defatted with alternate washes of a 2:1 (v/v) chloroform:methanol and 2:1 (v/v) methanol:chloroform solution in an ultrasonic bath for a maximum of 15 minutes each, until the solution remained clear. Another modification to the protocol described in Seiler et al. (2019) was the adjustment of temperature and concentration. Demineralisation was done in 0.2 to 0.5M HCl at 4°C until complete (24 hours to c. 4 weeks, depending on the sample particle size and preservation). Humic substances were removed with 0.1 M NaOH at 4ºC, left overnight, and repeated until the solution remained clear. Between/after reagents, samples were washed three times with ultrapure water until neutral. The samples were gelatinised at 70ºC and pH=3 for about 48 hours (or until complete), and freeze-dried. Combustion, graphitisation and measurement was performed at the National Laboratory for Age Determination (Trondheim, NO) as described in Seiler et al. (2019) and Nadeau et al. (2015). We used OxCal v.4.4.4 (Bronk Ramsey 2009; Bronk Ramsey and Lee 2013) to calculate ΔR values for the known-age specimens from natural history collections. Two alternative approaches were used. The first method used only date of death information, because it is known with considerable certainty. The second method included both date of death and inferred date of birth. Date of birth was estimated using the osteological age categories of Dierickx et al. (2025): neonatal (birth to 3/4 years), juvenile (c.4 to c.10 years) and adult (c.10 to c. 40 years); the midpoint of each age category was subtracted from date of death and attributed an error term such that two standard deviations would encompass the full age category. For adults, for example, 25 years was subtracted from date of death and the estimated date of birth then attributed an error of 7.5 (rounded to 8) years. Estimated date of birth was used to constrain the *terminus post quem* (TPQ) of the models because the bone collagen turnover rate for walruses is not well established.

The radiocarbon dates of these samples were placed in a simple sequence model, where the radiocarbon age (R_Date) was followed by the date of death (C_Date) (method 1) or bracketed by the estimated date of birth (C_Date) and the date of death (C_Date) (method 2). The uncertainty for date of death was set to ±1 year if known to a single year, or increased as appropriate in instances when harvest date was only known within several years. In the latter cases, additional years were also added to the error of the estimated date of birth. We calculated ΔRs per population. Furthermore, we calculated a pooled ΔR value for the populations that were potentially relevant for the medieval trade of walrus tusks, i.e., Greenland (excluding East Greenland) and the Canadian Arctic. There may be differences in diet and therefore in ΔR between individuals related to the size, age or sex of individuals, and specific feeding locations, but the aim of this study is to provide reliable ΔR estimates for specimens for which this information will typically not be available. Therefore, potential differences within populations are not a focus of this paper. As a final step, we used method 2 as described above to calculate ΔR values per population (and a pooled ΔR value for Greenland (excluding East Greenland) and the Canadian Arctic) when also including 15 radiocarbon dates on known-age walrus specimens pre-dating 1950 published by Dyke et al. (2019). One of these previously studied specimens (UCIAMS-168850) was re-examined within the present project (TRa-24150), but the 14 others were not examined in-hand for ontological age assessment. For the purposes of TPQ (C_Date) modelling, the date of birth of the Dyke et al. (2019) specimens was thus estimated as the date of death minus 20 years with a standard deviation of 10 years.

The twelve archaeological samples from Trondheim and its hinterland were calibrated three times. Firstly, we used the pooled ΔR calculated for the historical samples from Greenland and the Canadian Arctic, +17±12, and applied that to all samples. All OxCal code is provided in the Supplemental material. Secondly, we used the same ΔR but also included archaeological context information as described below.

For the third calibration of the archaeological samples, OxCal was used to estimate an appropriate ΔR in a similar way as for the historical specimens; as several samples originate from relatively well-dated contexts, we could use the date of the context in the same way as the year of death for historical samples (see e.g. Macario et al. 2023, van den Hurk et al. 2025). For example, if a sample’s context was dated to 1225–1275 CE, we assume a symmetrical distribution of the true calendar age around 1250 CE with ±25 years at 2σ. The C_Date for the model (with 1σ uncertainty) is thus 1250±12.5, rounded to 1250±13. In interpreting the results of this approach, it is necessary to account for the possibility that the year of incorporation into the archaeological record may be later than the year of death of the animal.

Table 4, Figures 5 and 6 display the results of these different approaches. All OxCal code can be found in online supplemental material 2, section 2.10.1 and 2.10.2.

**Table 4.**
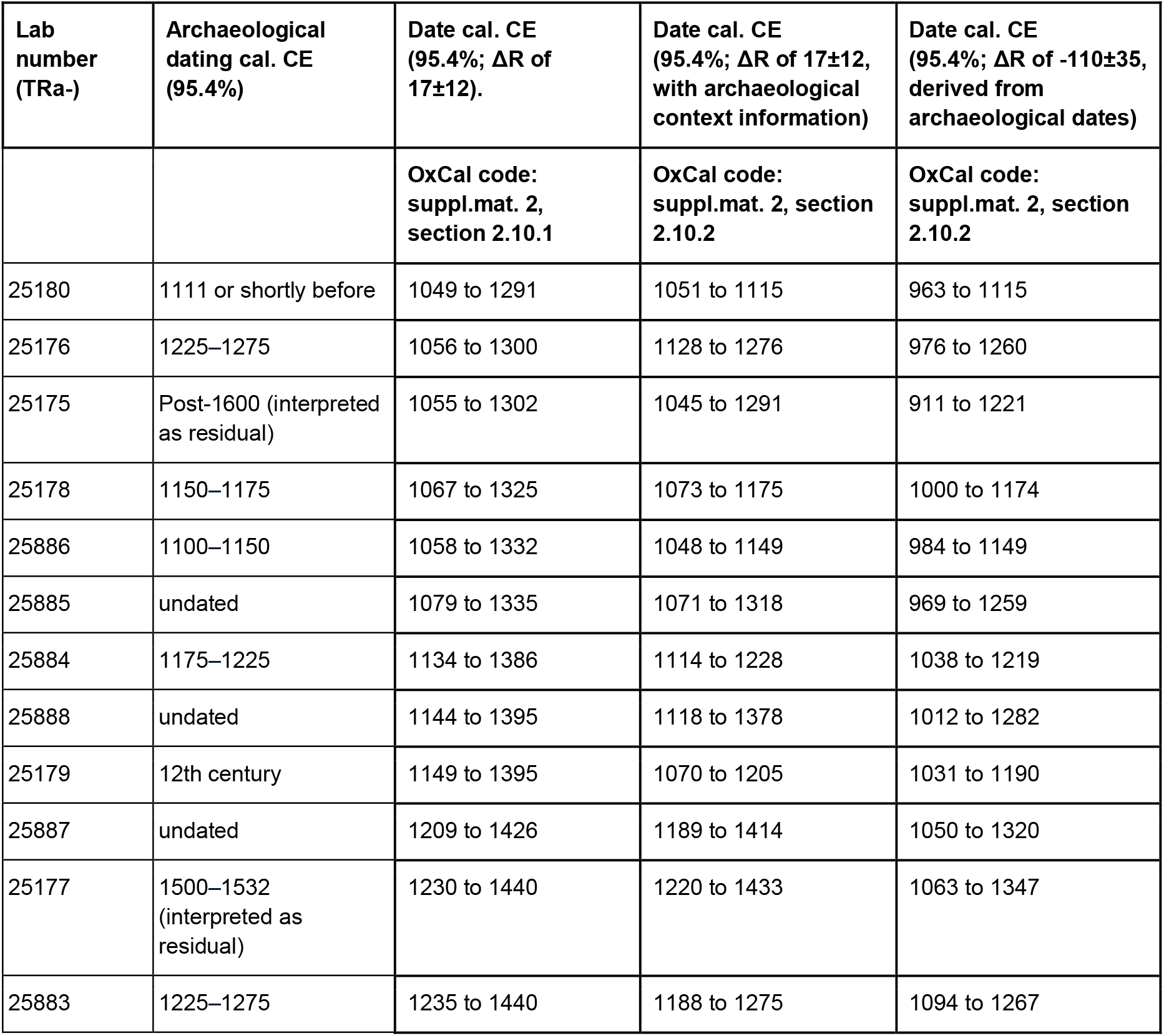
Calibrated date ranges CE (95.4%) for medieval walrus artefacts from Trondheim and its hinterland, calibrated in OxCal v4.4.4 (Bronk Ramsey 2009) using the Marine20 curve (Heaton et al. 2020) with three different approaches as described in the text. The OxCal code is provided in Supplemental material file 2 (section 2.10).

| Lab number (TRa-) | Archaeological dating cal. CE (95.4%) | Date cal. CE (95.4%; $\Delta R$ of $17 \pm 12$). | Date cal. CE (95.4%; $\Delta R$ of $17 \pm 12$, with archaeological context information) | Date cal. CE (95.4%; $\Delta R$ of $-110 \pm 35$, derived from archaeological dates) |
| --- | --- | --- | --- | --- |
|  |  | OxCal code: suppl.mat. 2, section 2.10.1 | OxCal code: suppl.mat. 2, section 2.10.2 | OxCal code: suppl.mat. 2, section 2.10.2 |
| 25180 | 1111 or shortly before | 1049 to 1291 | 1051 to 1115 | 963 to 1115 |
| 25176 | 1225–1275 | 1056 to 1300 | 1128 to 1276 | 976 to 1260 |
| 25175 | Post-1600 (interpreted as residual) | 1055 to 1302 | 1045 to 1291 | 911 to 1221 |
| 25178 | 1150–1175 | 1067 to 1325 | 1073 to 1175 | 1000 to 1174 |
| 25886 | 1100–1150 | 1058 to 1332 | 1048 to 1149 | 984 to 1149 |
| 25885 | undated | 1079 to 1335 | 1071 to 1318 | 969 to 1259 |
| 25884 | 1175–1225 | 1134 to 1386 | 1114 to 1228 | 1038 to 1219 |
| 25888 | undated | 1144 to 1395 | 1118 to 1378 | 1012 to 1282 |
| 25179 | 12th century | 1149 to 1395 | 1070 to 1205 | 1031 to 1190 |
| 25887 | undated | 1209 to 1426 | 1189 to 1414 | 1050 to 1320 |
| 25177 | 1500–1532 (interpreted as residual) | 1230 to 1440 | 1220 to 1433 | 1063 to 1347 |
| 25883 | 1225–1275 | 1235 to 1440 | 1188 to 1275 | 1094 to 1267 |

**Figure 5.**
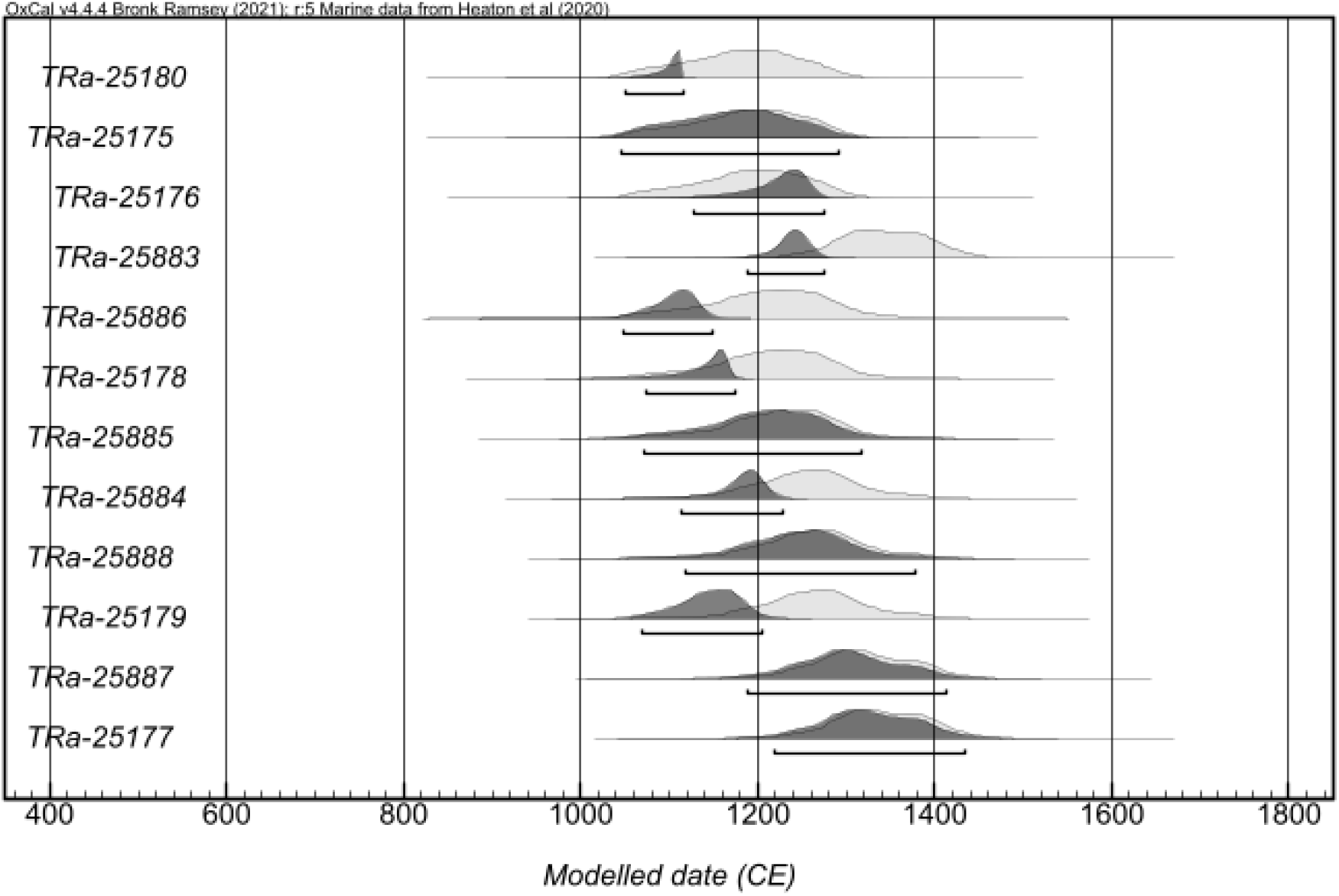
Calibrated ages of archaeological walrus samples from excavations in Trondheim and its hinterland, calibrated in OxCal v4.4.4 (Bronk Ramsey 2009) using the Marine20 curve (Heaton et al. 2020), the ΔR of 17±12 calculated for historical specimens from Greenland (excluding Eastern Greenland) and the Canadian Arctic, and prior information from the dates of the relevant archaeological contexts. See Table 2 for details and Table 4 for calibrated age ranges.

**Figure 6.**
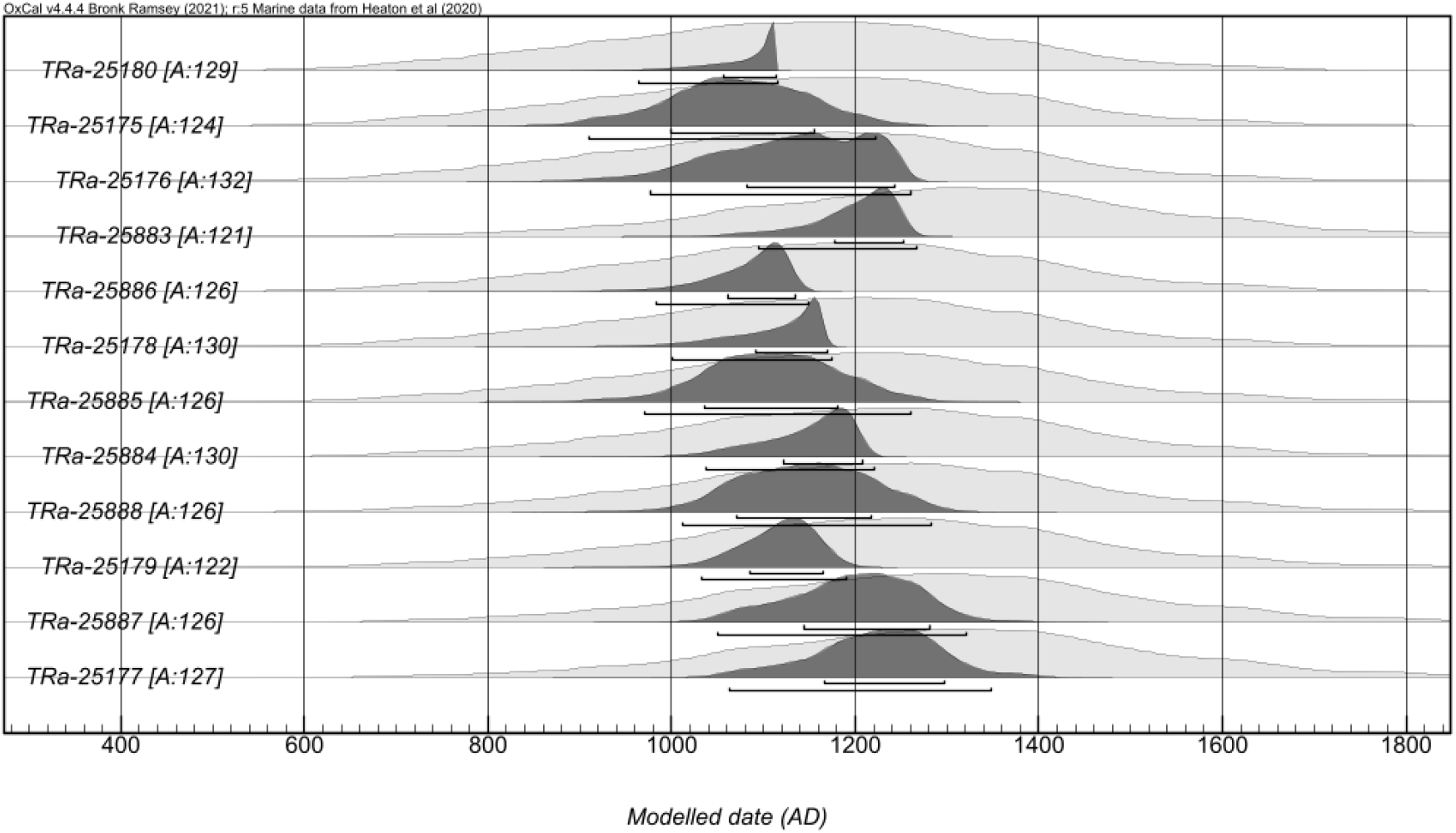
Alternative calibrated ages of archaeological walrus samples from excavations in Trondheim and its hinterland. For this figure, we did not specify a ΔR based on known-age natural history specimens, but let OxCal estimate the most appropriate ΔR based on the archaeological context dating, with a prior of U(−400,400). This resulted in ΔR between −184 and −40 (95.4%), with a median value of −107, mean −110, and standard deviation of 35. See also the last column of Table 4.

## Results and Discussion

### ΔR Values from Known-age Walrus Specimens

Based on the known-date natural history specimens, the calculated ΔR values per walrus population, and pooled for Greenland (excluding East Greenland) and the Canadian Arctic, are summarised in Table 3. The ΔR estimates calculated using the different methods agree within uncertainties (Table 3, Figure 4). Each estimate is valid within its own assumptions and limitations. Of the three approaches, the ΔR calculations including both estimated dates of birth and legacy dates are employed for the discussion that follows. Although these results entail more assumptions (regarding the age of the individual walruses) than the calculations using only date of death, they provide improved accuracy and precision.

There are considerable differences between the ΔR values for some populations. The mean ΔR values range from −140 for the samples from Franz Josef Land to +295 for the samples from the Pacific. The standard deviations also vary between populations; populations with a larger number of dated specimens have more restricted ΔR values.

The pooled ΔR for Greenland (excluding East Greenland) and the Canadian Arctic is +17±12. This value is heavily influenced by the samples from Baffin Bay (n=13) and Hudson Strait/Hudson Bay (n=14). The Baffin Bay samples alone have a ΔR of −12±19. This is only slightly lower than the value recorded in the 14CHRONO Marine20 Reservoir database at http://calib.org/marine/ (Reimer & Reimer 2001), drawing mainly on Pearce et al. (2023), where 17 shell samples from Baffin Bay have an average ΔR of +31±97. The 14 samples from the Hudson Strait and Hudson Bay provide a ΔR of +35±18, higher than the database estimate of −59±78 based on 35 shell and walrus samples, but within the 95.4% range. Western Greenland has a ΔR of −30±97 in the 14CHRONO Marine20 Reservoir database, based on 56 mostly shell samples (there is one seal and one walrus); this agrees closely with the ΔR of −17±47 which we calculated for the two walrus samples from that region. For the Foxe Basin, 44 shell samples and three walrus samples from the 14CHRONO Marine20 Reservoir database yield a ΔR of +164±122, higher than that derived from the five walrus samples considered here (ΔR = +49±29). It should be noted, however, that the range of ΔR from the Foxe Basin walrus samples is large (−10 to +109), overlapping with the database estimate. Our six walrus samples from Eastern Greenland have a ΔR value of −26±27, with a range from −82 to +29. In the reservoir database, 38 shell sample results are available for Eastern Greenland. These produce a ΔR of 84±244 but with a very large ΔR range, which overlaps with our results.

In the reservoir database, 14 shell samples from Iceland have ΔR = −94±76, which agrees well with our one walrus specimen from that island, which has ΔR = −49±66. However, as this is only one sample, its range of possible ΔR values is broad (from −182 to +84). It should also be noted that the Icelandic walrus population is thought to have been extirpated in the Viking Age (Keighley et al. 2019); the individual we dated (killed in 1900) was probably a vagrant from another population, having unknown residency time in Icelandic waters.

From Franz Josef Land, two shell measurements are available in the reservoir database (note that sample GX-19024, from the bivalve *Bathyarca glacialis*, is mis-attributed to whale in the database, see Pieńkowski et al. 2022). Both have negative ΔR values, −283 and −265, producing an average ΔR of −276±31 (see also Pieńkowski et al. 2022). This is lower than the ΔR calculated for our two walrus samples from Franz Josef Land, −140±47, but within the 95.4% range. It may be pertinent that the mollusk-derived ΔR for Svalbard (−61±37) is also higher than that from Franz Josef Land (Pieńkowski et al 2022). Some migration of walruses is documented between the two archipelagoes (e.g. Mikkelsen et al. 2024), so walrus ΔR values in the Barents Sea may vary depending on the degree of feeding in each group of islands.

The exceptionally high ΔR value of our two samples from the Pacific (295±47) is supported by 12 shell samples from the reservoir database, with ΔR= 305±102. Our Pacific value is also comparable with the published ΔR of 289±124 based on terrestrial versus marine (seal and walrus) sample pairs from archaeological contexts in coastal Chukotka and Alaska (Dury et al. 2021).

Overall, the results based on known-age samples indicate the importance of using population-specific ΔR values for walruses, especially in extreme cases such as the Pacific and Franz Josef Land, but to a lesser degree also in other areas. Where both shell- and walrus-derived ΔR values exist, species and population-specific walrus ΔR values are preferable. However, most walrus and shell data are broadly consistent, justifying use of shells where known-age walrus specimens do not exist. This would apply, for example, when calibrating radiocarbon dates for the Maritimes walrus of eastern Canada, which was extirpated in the 18th century CE, leaving no extant known-age specimens (McLeod et al. 2014; McCaffrey 2020). Caution does remain necessary in locations where there are few or no known-date walrus samples and shell data indicate high intra-regional variability (e.g. East Greenland). The seasonal mobility of walruses should dampen some local variability in shell-based ΔR values, but high feeding-site fidelity (e.g. Mikkelsen et al. 2024) means that ΔR variability within walrus populations could nevertheless occur where baseline food-source values are heterogeneous.

### Calibration of Radiocarbon Dates on Archaeological Walrus Finds from the Trondheim Region

The walrus rostra and ivory finds from archaeological contexts in Trondheim and its hinterland are presently interpreted as evidence of trade from the medieval Norse colonies of Greenland (Barrett et al. 2020; Ruiz-Puerta et al. 2024). This interpretation is based on aDNA and/or stable isotope provenancing of a subset of the relevant finds (including six of the 12 specimens that are radiocarbon dated in this paper: N10102, N37603, N32091, N29494, N203145 and N167587), showing that they probably derived from western/northwestern Greenland or the eastern Canadian Arctic. Yet two of the provenanced specimens (N37603 and N167587) are from archaeological contexts that postdate the traditionally accepted 15th-century CE abandonment of Norse Greenland and pre-date the subsequent Scandinavian return to the island in the 18th century (Christophersen and Nordeide 1994; Nordeide 2020; Gulløv 2016).The question thus arises whether these are earlier objects that have been redeposited in later archaeological layers (i.e. they are residual), or if Norwegian trade with Greenland continued later than assumed.

To address this question, we used our pooled ΔR values for Greenland (excluding East Greenland) and the Canadian Arctic (17±12) to calibrate the radiocarbon dates of these finds, plus 10 other walrus artefacts from Trondheim (three unstratified and seven from 12th and 13th-century contexts) (Table 2, Figure 5). In the first instance, we calibrated the twelve archaeological dates as a single group without stratigraphic information (Table 4). In a second OxCal calibration, we added the original archaeological context dating (except for the two potentially residual specimens, which were treated as unstratified) in a simple Bayesian model. In cases where the archaeological date of the layer in which the sample was found is known, the walrus must have been killed before the deposition in this layer, and we used simple sequence models as for the historical specimens. Where no archaeological dating is available, the relevant cell in Table 2 is left blank and no sequence model was used for that sample. In result, there is variable agreement between the walrus radiocarbon and archaeological dates; the agreement indices (A) are below the recommended threshold value of 60 for five specimens: TRa-25883, TRa-25886, TRa-25178, TRa-25884 and TRa-25179, suggesting that some may be intrusive or residual rather than being in their primary depositional context. Less likely, some could alternatively derive from walruses having a different ΔR (see further below). Regardless, the calibrated results from both approaches are consistent with all 12 objects having derived from between the 11th and early 15th centuries (Table 4). This dating corresponds with the known medieval trade in walrus ivory between Greenland and Norway (Barrett 2021; Barrett and Grav 2025) and is before the mid-15th-century abandonment of Norse Greenland (Arneborg 2024).

In a third calibration we excluded the predetermined ΔR of 17±12, and instead employed code allowing OxCal to calculate ΔR using only the archaeological context dates for reference. The result is −110±35 (Figure 6), lower than the ΔR calculated for the historical samples. At face value, the negative ΔR derived from the archaeological dating could indicate a smaller marine reservoir offset than we have calculated using historical specimens. However, the difference from a ΔR of 17±12 is more likely to reflect the elapsed time between when the walruses were harvested and when the resulting raw materials reached their final archaeological context (after transport, storage, use, primary deposition and potentially secondary or tertiary redeposition). If so, the ΔR based on the known-age natural history specimens will be more reliable. Alternatively, a ΔR difference between 19th- to 20th-century and medieval walruses from the same region could indicate changes in feeding behaviour. This possibility can be explored by stable isotope (δ^13^C, δ^15^N, δ^34^S) analysis in future research (Dierickx et al. forthcoming). If the Trondheim finds are calibrated assuming a ΔR of −110±35 (Table 4), the resulting date ranges are slightly older than when 17±12 was employed (given the smaller marine reservoir adjustment of the raw radiocarbon determinations). However, regardless of which of the three calibration approaches is adopted, it is clear that all of the specimens dated relate to the centuries during which walrus ivory trade between the Norse colonies of Greenland and Norway is plausible based on existing archaeological and historical evidence (Arneborg 2021).

## Conclusion

The large differences in ΔR values for the different populations show that it is important to consider the population of origin when radiocarbon dating walrus samples. This can be a challenge for samples of unknown provenance (e.g., archaeological samples). Provenance studies (e.g., using aDNA and stable isotopes) may thus be a prerequisite for accurate radiocarbon dating of walrus samples. In many instances, walrus and shell samples from the same localities produce similar ΔR values, indicating that shell data can often serve as an acceptable alternative where known-age walrus specimens do not exist, as is the case for extirpated populations. Caution is required, however, in regions with heterogeneous ΔR values based on shell or walrus samples. Using our ΔR results from known-date natural history specimens to calibrate radiocarbon dates on walrus artefacts from Trondheim it is possible to confirm that seemingly anomalous archaeological dates are due to redeposition of residual artefacts.

## Supporting information

Supplemental Material 2: OxCal codes

Supplemental Material 3: Data from calib.org-marine

## Acknowledgments

We thank the following institutions and individuals for providing specimens and documentation: the American Museum of Natural History (New York, USA): Marisa Surovy; Canadian Museum of Nature (Ottawa and Gatineau, Canada): Scott Rufolo, Shyong En Pan, Gregory Rand; Natural History Museum (London, United Kingdom): Roberto Portela Miguez, Phaedra Kokkini; Zoological Museum, Natural History Museum of Denmark: Peter Rask Møller, Daniel Klingberg Johansson; NTNU University Museum: Birgitte Skar, Thea P. B. Christophersen, Terje Masterud Hellan, Torkel Johansen, Birgit Maixner, Jon Anders Risvaag, Marte Iversen Rønning. Magnar Mojaren Gran kindly assisted with Figure 1. Axel Christophersen was generous with his insights regarding the context of the Trondheim finds. Martin Seiler assisted with the radiocarbon dating. Bastiaan Star, Lydia Furness and Oliver Kersten assisted with the subsampling of specimens also destined for ancient DNA analysis. The Board of Directors and Office Manager of the Igloolik Hunter’s and Trapper’s Association kindly provided feedback on the results from an Inuit perspective.

## Ethical approval and informed consent statements

The walrus specimens studied herein are all from registered museums. No live animals were collected for the research and no fieldwork was conducted. Sampling permissions for natural history specimens were kindly provided by the American Museum of Natural History, the Canadian Museum of Nature, the Natural History Museum (London) and the Zoological Museum, Natural History Museum of Denmark. Sampling permission for the Norwegian archaeological specimens was kindly agreed by the NTNU University Museum. Samples were shipped across borders (from Canada, Denmark, the United Kingdom, and United States of America to Norway) under institutional CITES exemptions CA 003, DK003, GB 001, NO 001, NO 007, and US 055 (permit number 24US66924E/9).

## Data availability statement

All raw data can be found in supplemental material file 1, containing Tables S1 and S2. OxCal code is provided in supplemental material file 2.

## List of Supplemental Material

- Supplemental Material 1.
  - Table S1. Metadata for radiocarbon dated walrus samples with historical dates of death. European place-names are retained if needed to avoid ambiguity.
  - Table S2. Metadata for radiocarbon dated walrus specimens from excavations in Trondheim and its hinterland.
  - Table S3. Pre-1950 radiocarbon dated walrus samples with historical dates of death from Dyke et al. (2019).
- Supplemental Material 2. OxCal code for ΔR determinations by population or location and for calibration of the archaeological specimens from Trondheim and its hinterland
- Supplemental Material 3. Predominantly shell-derived ΔR values from areas relevant to the walrus populations considered in this paper, from the 14CHRONO Marine20 Reservoir database at http://calib.org/marine/ (Reimer & Reimer 2001).

