## Supplemental Material 1: Tables for "Walrus Population-specific Marine Reservoir Offsets (ΔR) for Calibration of Radiocarbon Dates: Implications for Arctic Chronologies and Medieval Trade"

**Table S1. Metadata for radiocarbon dated walrus samples with historical dates of death. European place-names are retained if needed to avoid ambiguity.**

**Table S2. Metadata for radiocarbon dated walrus specimens from excavations in Trondheim and its hinterland.**

**Table S3. Pre-1950 radiocarbon dated walrus samples with historical dates of death from Dyke et al. (2019).**

**Table S1. Metadata for radiocarbon dated walrus samples with historical dates of death. European place-names are retained if needed to avoid ambiguity.**

| Lab number | 4-OCEANS unique identifier | Estimated age following Dierickx et al. (2025) | Source | Museum catalogue number | Population | Origin as historically catalogued | Origin | Country | Tissue sampled | Skeletal element | Location on element | Decimal lat. | Decimal long. | Date of death | Date first registered | Explanation of historical dating | Unrounded age BP | Unrounded error | General comments |
|---|---|---|---|---|---|---|---|---|---|---|---|---|---|---|---|---|---|---|---|
| TRa-22945 | 3829.1 | adult | Natural History Museum, London | NHMUK ZD.1890.12.4.2 | Baffin Bay | Wellington Channel Polar Seas | Wellington Channel | Canada | bone | cranium | mastoid | 75.196 | -92.97 | 1852 to 1853 | 1890 | date of collection must be Robert McCormick's 1852 to 1853 Canadian expedition in search of the missing Franklin Expedition; pres. Dr R. <on label> or P <in catalogue> McCormick R.N.; Wellington Channel Arctic Seas | 649 | 12 | pres. Dr R. <on label> or P <in catalogue> McCormick R.N.; Wellington Channel Arctic Seas; complete skull with tusks in place; mandible attached to skull; occipital condyles cut through in transverse plane from decapitation |
| TRa-22943 | 3827.1 | adult | | NHMUK ZD.1855.11.26.38 | Baffin Bay | Barrow Straits | Barrow Strait | Canada | bone | cranium | tympanic bulla | 74.481 | -97.196 | 1853 to 1854 | 1855 | observation that Inglefield was captain of HMS Phoenix and undertook two voyages to Arctic during 1853-1854 | 725 | 15 | collected by Capt. Inglefield; H.M.S. Phoenix; from Barrow Straits; information on label is also written in decorative lettering on front of rostrum |
| TRa-22944 | 3828.1 | adult | | NHMUK ZD.1855.11.26.39 | Baffin Bay | Barrow Straits | Barrow Strait | Canada | bone | cranium | maxilla | 74.481 | -97.196 | 1854 | 1855 | pres. to Mr Barrow by Capt Jenkins H.M.S. Talbot; Barrow Straits; information on label (plus the year 1854) is also written in decorative lettering on the back of the left tusk | 737 | 13 | pres. to Mr Barrow by Capt Jenkins H.M.S. Talbot; Barrow Straits; information on label (plus the year 1854) is also written in decorative lettering on the back of the left tusk; rostrum (severed from skull by chopping from dorsal) with tusks in place |
| TRa-25066 | 4859.1 | adult | American Museum of Natural History | M-14068 | Baffin Bay | Murchison Sound | Murchison Sund | Greenland (Denmark) | bone | cranium | tympanic bulla | 77.333 | -71.5 | 1895 | 1895 | museum dates as 1 Oct 1895; Dyche was on expedition some time between 1894 and 1895 | 582 | 13 | collected by L.L. Dyche |
| TRa-25063 | 4853.1 | adult | | M-14071 | Baffin Bay | Murchison Sound | Murchison Sund | Greenland (Denmark) | bone | cranium | tympanic bulla | 77.333 | -71.5 | 1896 | 1896 | museum dates as Oct 1896 which is plausible given Peary's trips to Greenland in 1895 and 1896 | 569 | 21 | collected by Robert Peary |
| TRa-25064 | 4855.1 | adult | | M-14072 | Baffin Bay | Murchison Sound | Murchison Sund | Greenland (Denmark) | bone | cranium | temporal | 77.333 | -71.5 | 1896 | 1896 | museum dates as 1 Oct 1896 which is plausible given Peary's trips to Greenland in 1895 and 1896 | 535 | 13 | collected by Robert Peary |
| TRa-25065 | 4857.1 | adult | | MO-11049 | Baffin Bay | Manson Island | Qeqertaarsuit (Manson Øer) | Greenland (Denmark) | bone | cranium | tympanic bulla | 76.654 | -69.17 | 1896 | 1896 | museum dates as 20 August 1896 which is plausible given Peary's trips to Greenland in 1895 and 1896 | 599 | 15 | collected by Robert Peary |
| TRa-25068 | 4874.1 | adult | | MO-11045 | Baffin Bay | Manson Island | Qeqertaarsuit (Manson Øer) | Greenland (Denmark) | bone | cranium | temporal | 76.654 | -69.17 | 1896 | 1896 | museum dates as 11 Aug 1896 which is plausible given Peary's trips to Greenland in 1895 and 1896 | 603 | 14 | collected by Robert Peary |
| TRa-25069 | 4875.1 | adult | | MO-11046 | Baffin Bay | Manson Island | Qeqertaarsuit (Manson Øer) | Greenland (Denmark) | bone | cranium | tympanic bulla | 76.654 | -69.17 | 1896 | 1896 | museum dates as 11 Aug 1896 which is plausible given Peary's trips to Greenland in 1895 and 1896 | 582 | 13 | collected by Robert Peary |
| TRa-25060 | 4849.1 | adult | | M-73301 | Baffin Bay | Northumberland Island, Greenland | Kiatak | Greenland (Denmark) | bone | cranium | temporal (zygomatic process) | 77.383 | -71.933 | 1926 | 1926 | museum dates as 17.08.1926; Raven was on expedition in 1926 to Greenland for museum | 553 | 16 | collected by H.C. Raven |
| TRa-25061 | 4851.1 | juvenile | | M-73303 | Baffin Bay | Northumberland Island, Greenland | Kiatak | Greenland (Denmark) | bone | cranium | tympanic bulla | 77.383 | -71.933 | 1926 | 1926 | museum dates as 17.08.1926; Raven was on expedition in 1926 to Greenland for museum | 563 | 13 | collected by H.C. Raven |
| TRa-25062 | 4852.1 | adult | | M-73304 | Baffin Bay | Northumberland Island, Greenland | Kiatak | Greenland (Denmark) | bone | cranium | parietal | 77.383 | -71.933 | 1926 | 1926 | museum dates as 17.08.1926; Raven was on expedition in 1926 to Greenland for museum | 565 | 18 | collected by H.C. Raven |
| TRa-25067 | 4860.1 | juvenile | | MO-10176 | Western Greenland | Holsteinborg (Sisimiut) | Sisimiut | Greenland (Denmark) | bone | cranium | tympanic bulla | 66.941 | -53.674 | 1895 | 1895 | museum dates as 8 Oct 1895; Dyche was on expedition some time between 1894 and 1895 | 611 | 20 | collected by L.L. Dyche |
| TRa-25591 | 4684.1 | adult | Zoological Museum, Natural History Museum of Denmark, Copenhagen | NHMD-M11-CN276 | Western Greenland | Holsteinborg | Sisimiut | Greenland (Denmark) | bone | cranium | tympanic bulla | 66.941 | -53.674 | 1898 | 1899 | killed 12-3-1898; registered 24.05.1899 | 604 | 13 | collected by colony manager R. Müller |
| TRa-25595 | 4751.1 | juvenile | | NHMD-M11-CN480 | cf. Foxe Basin | Frozen Strait (just north of Hudson Bay) | Frozen Strait | Canada | bone | cranium | tympanic bulla | 65.845 | -84.44 | 1922 | 1922 | killed Dec 1922; collected by Peter Freuchen on the 5th Thule expedition | 646 | 18 | collected by Peter Freuchen on the 5th Thule expedition |
| TRa-25596 | 4752.1 | juvenile | | NHMD-M11-CN479 | cf. Foxe Basin | Frozen Strait (just north of Hudson Bay) | Frozen Strait | Canada | bone | cranium | tympanic bulla | 65.845 | -84.44 | 1922 | 1922 | killed Dec 1922; collected by Peter Freuchen on the 5th Thule expedition | 687 | 16 | cutmarks; collected by Peter Freuchen on the 5th Thule expedition |
| TRa-24149 | 4512.1 | juvenile | Canadian Museum of Nature | CMNMA 37 | Hudson Strait | Nunavut; Nottingham Island | Tujjaat | Canada | bone | cranium | tympanic bulla | 63.349 | -77.985 | 1884 to 1885 | 1885 | first or second Hudson Bay Expedition by Gordon in 1884/1885 that passed locality; Bell was on board as medical officer | 671 | 20 | collected by Dr. R. Bell |
| TRa-24150 | 4516.1 | juvenile | | CMNMA 36 | Hudson Strait | Prince of Wales Sound; Hudson Strait | Hudson Strait | Canada | bone | cranium | temporal (squamosal) | 61.562 | -71.583 | 1884 to 1885 | 1885 | first or second Hudson Bay Expedition by Gordon in 1884/1885 that passed locality; Bell was on board as medical officer | 639 | 13 | collected by Dr. R. Bell; location probably refers to an old name on Hudson Strait in modern Nunavut not Prince of Wales Strait in what is now NWT; See https://recherche-collection-search.bac-lac.gc.ca/eng/Home/Record?app=fonandcol&IdNumber=3328314&ecopy= |
| TRa-24148 | 4511.1 | adult | | CMNMA 93 | Hudson Strait | Ashe Inlet Hudson Strait | Ashe Inlet | Canada | bone | mandible | tooth socket (most distal) | 62.546 | -70.558 | 1886 | 1886 | date of death specifically given as 1886-04-10 (10 April if normal Canadian date format) in email from Gregory Rand on 2 April 2025 | 609 | 12 | some other teeth have been glued in but no tooth or glue in the small distal socket sampled; museum catalogue records 11' 0" and 2500 lb |
| TRa-24151 | 4519.1 | adult | | CMNMA 1898 | cf. Hudson Strait | Fort Churchill; Manitoba; Hudson Bay | Churchill | Canada | bone | cranium | tusk socket fragment near lip on palatal side | 58.757 | -94.083 | 1910 | 1910 | Thomas N. Marcellus was present in Fort Churchill in 1910 | 615 | 12 | collected by Dr. Marcellus |
| TRa-25587 | 4676.1 | adult | Zoological Museum, Natural History Museum of Denmark, Copenhagen | NHMD-M11-CN732 | Eastern Greenland | Scoresby Sund | Kangertittivaq | Greenland (Denmark) | bone | cranium | tympanic bulla | 70.613 | -24.701 | 1927 | 1927 | this time and regularly sent his collection to museum; his trip was from 1927-29 | 605 | 23 | collected by Alwin Pedersen |
| TRa-25588 | 4679.1 | adult | | NHMD-M11-CN729 | Eastern Greenland | Scoresby Sund | Kangertittivaq | Greenland (Denmark) | bone | cranium | tympanic bulla | 70.613 | -24.701 | 1927 to 1929 | 1935 | Scoresby Sund and regurlarly sent his collection to museum; his trip was from 1927-29 | 558 | 16 | collected by Alwin Pedersen |
| TRa-25585 | 4671.1 | adult | | NHMD-M11-CN726 | Eastern Greenland | Scoresby Sund | Kangertittivaq | Greenland (Denmark) | bone | cranium | tympanic bulla | 70.613 | -24.701 | 1929 | 1929 | Sund at this time and regurlarly sent his collection to museum; his trip was from 1927-29 | 561 | 16 | collected by Alwin Pedersen |
| TRa-25586 | 4674.1 | juvenile | | NHMD-M11-CN727 | Eastern Greenland | Scoresby Sund | Kangertittivaq | Greenland (Denmark) | bone | cranium | tympanic bulla | 70.613 | -24.701 | 1929 | 1929 | museum record gives 25/09/1929; Alwin Pedersen was in Scoresby Sund at this time and regurlarly sent his collection to museum; his trip was from 1927-29 | 567 | 15 | collected by Alwin Pedersen |
| TRa-25589 | 4682.1 | adult | | NHMD-M11-CN293 | Eastern Greenland | North east Greenland | Tunu | Greenland (Denmark) | bone | cranium | maxilla posterior to cheek teeth | 80.329 | -18.237 | 1906 to 1909 | 1909 | registered 18.03.1909; F. Johansen was a member of Danish expedition in 1906-1908 to Greenland's north-east coast led by Mylius-Erichsen | 631 | 21 | collected by F. Johansen |
| TRa-25593 | 4749.1 | adult | | NHMD-M11-CN461 | Eastern Greenland | Scoresby Sund | Kangertittivaq | Greenland (Denmark) | bone | cranium | tympanic bulla | 70.613 | -24.701 | 1924 | 1924 | museum label records as killed August 1924 and registered 14 November 1924; collected by Alwin Pedersen | 569 | 13 | collected by Alwin Pedersen |
| TRa-25590 | 4683.1 | adult | | NHMD-M11-CN273 | cf. Iceland | Thorlakshofn Iceland | Þorlákshöfn | Iceland | bone | mandible | tooth socket; 1st mesial | 63.86 | -21.404 | 1900 | 1905 | registered 14.05.1905; shot in spring 1900 by a farmer named Jon Arnason | 576 | 14 | shot by Jon Arnason |
| TRa-22947 | 3831.1 | adult | Natural History Museum, London | NHMUK ZD.1929.7.24.10 | Franz Josef Land | Franz Josef Land | Franz Josef Land | Russia | bone | mandible | tooth socket | 80.805 | 54.925 | 1894 to 1897 | 1929 | observation that Jackson-Harmsworth expedition was led by F.G. Jackson between 1894 and 1897 | 494 | 15 | pres. Major F.G. Jackson; Franz Josef Land |
| TRa-22952 | 3837.1 | neonatal | | NHMUK ZD.1897.10.22.1 | Franz Josef Land | Cape Flora Franz Josef Land | Cape Flora | Russia | bone | cranium | tympanic bulla | 79.948 | 50.102 | 1894 to 1897 | 1897 | date first registered inferred from catalogue number; date of death based on observation that Jackson-Harmsworth expedition led by F.G. Jackson between 1894-1897 | 472 | 15 | pres. F.G. Jackson Esq.; Cape Flora Franz Josef Land; complete neonatal skull with tusk buds and separate mandible; some cranial sutures are glued but no treatment where sampled |
| TRa-22951 | 3836.1 | juvenile | | NHMUK GERM.331j | Pacific | Arctic Ocean; 75°10' N; 168°24' W | Arctic Ocean | Arctic Ocean | bone | cranium | tympanic bulla | 75.221 | 168.45 | 1865 | late 19th-century | degrees 24 minutes W Long; Aug 21 1865; ship built 1852 bought by Towns in 1864; lost 1873; details in (original) decorative lettering on front of left tusk socket | 843 | 13 | captured by barque R. Towns; Arctic Ocean; 75 degrees 10 minutes N Lat; 168 degrees 24 minutes W Long; Aug 21 1865; all catch details written in decorative lettering (framed by careful scrollwork) on front of tusk sockets |
| TRa-22950 | 3834.1 | adult | | NHMUK ZD.1903.1.25.1 | Pacific | Kamtchatka | Kamchatka | Russia | bone | cranium | tympanic bulla | 56.203 | 159.7 | 1896 to 1897 | 1898 | 1903 Mammals register states 'received in 1898'; date of death based on observation that Thompson was on expedition in 1896-1897 to Bering Strait | 1026 | 16 | pres. Prof. D'Arcy W. Thompson; “Kamtchatka” |

**Table S2. Metadata for radiocarbon dated walrus specimens from excavations in Trondheim and its hinterland.**

| Lab number | 4-OCEANS unique identifier | Source | Museum number | Archaeological site | Context | Description | Sample material | Lat. | Long. | Archaeological context date (CE) | Context interpretation | Unrounded age BP | Unrounded error | Rostrum number (Barrett et al. 2020) |
|---|---|---|---|---|---|---|---|---|---|---|---|---|---|---|
| TRa-25177 | 4812.1 | NTNU University Museum | N167587 | Erkebispegården | H564; Felt 91/14; Lag (542); 14/6/95; period 6, phase 2, Group 156 | rostrum | bone | 63.4258 | 10.396 | 1500-1532 | residual? | 1229 | 13 | R1 |
| TRa-25175 | 4810.1 | | N37603 | Folkebibliotekstomten | FA (592); 26/05/76; phase 12; delfase 9 in Meddelelser 4 | rostrum | bone | 63.4309 | 10.4014 | post 1600 | residual? | 1389 | 21 | R4 |
| TRa-25176 | 4811.1 | | N32091 | Folkebibliotekstomten | FL (359); 11/8/75; phase 8; delfase 9 in Meddelelser 9 | rostrum fragment | bone | 63.4311 | 10.4014 | 1225-1275 | in situ | 1388 | 14 | R3 |
| TRa-25178 | 4813.1 | | N29494 | Folkebibliotekstomten | FA (436); 8/7/75; phase 6; delfase 3 in Meddelelser 4 | rostrum | root of cheek tooth | 63.4309 | 10.4014 | 1150-1175 | in situ | 1363 | 20 | R2 |
| TRa-25883 | 5358.1 | | N19934 | Folkebibliotekstomten | FA (245); 20/7/74; phase 8; delfase 5 in Meddelelser 4 | tusk offcut | tusk | 63.4309 | 10.4014 | 1225-1275 | in situ | 1226 | 13 | |
| TRa-25884 | 5359.1 | | N30109a | Folkebibliotekstomten | FK(463); 19/7/75; phase 7; delfase 7 in Meddelelser 5 | tusk offcut | tusk | 63.4312 | 10.4013 | 1175-1225 | in situ | 1321 | 12 | |
| TRa-25886 | 5362.1 | | N33186 | Folkebibliotekstomten | FF (983); 1/9/75; phase 5; delfase 5 in Meddelelser 3 | tusk gaming piece | tusk | 63.4311 | 10.4012 | 1100-1150 | in situ | 1364 | 34 | |
| TRa-25887 | 5363.1 | | T4869 | Kjøpmannsgata | donation of 1896; originally found during digging on Kjøpmannsgata near the old city bridge (den gamle bybro); thought likely to derive from Dominican monastery | tusk crucifix corpus | tusk | 63.4284 | 10.3999 | undated | unstratified | 1255 | 11 | |
| TRa-25885 | 5360.1 | | N41911 | Kjøpmannsgata 25-27 | KG (19); 30/3/77 | tusk offcut | tusk | 63.4304 | 10.4022 | undated | unstratified | 1351 | 12 | |
| TRa-25179 | 5297.1 | | N203145 | Kjøpmannsgata Waterfront | Felt 93/2 (2699) | rostrum fragment | bone | 63.4296 | 10.4014 | 12th century | in situ | 1312 | 13 | R5 |
| TRa-25888 | 5364.1 | | T2383:2 | Rømesfjeldet, Rømmen, Åfjorden | Rømesfjeldet, Rømen, Åfjorden | tusk (complete) | tusk | 63.983 | 10.4762 | undated | unstratified | 1313 | 14 | |
| TRa-25180 | 5298.1 | | N10102 | Søndre gate | Felt II, Felt T (103) | rostrum fragment | bone | 63.4306 | 10.4001 | 1111 or shortly before | in situ | 1399 | 16 | R6 |

**Table S3. Pre-1950 radiocarbon dated walrus samples with historical dates of death from Dyke et al. (2019).**
**Decimal degree georeferences and locality extracted from the 14CHRONO Marine20 Reservoir database at http://calib.org/marine/ (Reimer & Reimer 2001)**

| Lab number | Canadian Museum of Nature number | Matching sample TRa lab number | Matching sample 4-OCEANS unique identifier | Latitude | Longitude | Population as defined in the present paper | Reference | Locality | Date of death | Reported age BP | Reported error |
|---|---|---|---|---|---|---|---|---|---|---|---|
| UCIAMS-168854 | 5968 | NA | NA | 76.2092 | -81.019 | Baffin Bay | Dyke et al. 2019 | Craig Harbour, Ellesmere Island | 1924 | 755 | 15 |
| UCIAMS-168833 | 21740 | NA | NA | 67.8622 | -77.2033 | Foxe Basin | Dyke et al. 2019 | Foxe Basin | 1949 | 685 | 15 |
| UCIAMS-168834 | 21741 | NA | NA | 67.8622 | -77.2033 | Foxe Basin | Dyke et al. 2019 | Foxe Basin | 1949 | 655 | 15 |
| UCIAMS-168835 | 21742 | NA | NA | 67.8622 | -77.2033 | Foxe Basin | Dyke et al. 2019 | Foxe Basin | 1949 | 595 | 15 |
| UCIAMS-168836 | 21960 | NA | NA | 61.5972 | -71.962 | Hudson Strait | Dyke et al. 2019 | Central Hudson Strait | 1885 | 670 | 15 |
| UCIAMS-168850 | 36 | TRa-24150 | 4516.1 | 61.5972 | -71.962 | Hudson Strait | Dyke et al. 2019 | Central Hudson Strait | 1885 | 665 | 15 |
| UCIAMS-168857 | 10429 | NA | NA | 62.5512 | -70.595 | Hudson Strait | Dyke et al. 2019 | Central Hudson Strait | 1886 | 630 | 15 |
| UCIAMS-168851 | 32372 | NA | NA | 62.9882 | -82.2433 | Hudson Strait | Dyke et al. 2019 | Western Hudson Strait & Bay | 1923 | 595 | 15 |
| UCIAMS-168852 | 5548 | NA | NA | 63.676 | -77.3712 | Hudson Strait | Dyke et al. 2019 | Western Hudson Strait & Bay | 1924 | 660 | 15 |
| UCIAMS-168853 | 5549 | NA | NA | 63.676 | -77.3712 | Hudson Strait | Dyke et al. 2019 | Western Hudson Strait & Bay | 1924 | 690 | 15 |
| UCIAMS-168855 | 5983 | NA | NA | 63.676 | -77.3712 | Hudson Strait | Dyke et al. 2019 | Western Hudson Strait & Bay | 1924 | 670 | 15 |
| UCIAMS-168856 | 10428 | NA | NA | 63.676 | -77.3712 | Hudson Strait | Dyke et al. 2019 | Western Hudson Strait & Bay | 1924 | 695 | 15 |
| UCIAMS-168831 | 10368 | NA | NA | 64.2315 | -76.5422 | Hudson Strait | Dyke et al. 2019 | Western Hudson Strait & Bay | 1928 | 705 | 15 |
| UCIAMS-185717 | 10353 | NA | NA | 64.2315 | -76.5422 | Hudson Strait | Dyke et al. 2019 | Western Hudson Strait & Bay | 1928 | 625 | 15 |
| UCIAMS-168832 | 19323 | NA | NA | 64.2315 | -76.5422 | Hudson Strait | Dyke et al. 2019 | Western Hudson Strait & Bay | 1945 | 635 | 15 |
