## Supplemental Material 2: OxCal codes for "Walrus Population-specific Marine Reservoir Offsets (ΔR) for Calibration of Radiocarbon Dates: Implications for Arctic Chronologies and Medieval Trade"

**Supplemental Material 2. OxCal code for ΔR determinations by population or location and for calibration of the archaeological specimens from Trondheim and its hinterland**

**2.1.1 Baffin Bay, including date of death only**

```
Plot()
 {
  Curve("Marine20", "marine20.14c");
  Delta_R("BAF",U(-500,500));
  Sequence("Year of death 1853")
  {
   Boundary("Start 1853");
   R_Date("TRa-22945",649,12);
   C_Date("1853",1853,2);
   Boundary("End 1853");
  };
  Sequence("Year of death 1854")
  {
   Boundary("Start 1854");
   Phase("1854 walrus")
   {
    R_Date("TRa-22943",725,15);
    R_Date("TRa-22944",737,13);
   };
   C_Date("1854",1854,2);
   Boundary("End 1854");
  };
  Sequence("Year of death 1895")
  {
   Boundary("Start 1895");
   R_Date("TRa-25066",582,13);
   C_Date("1895",1895,1);
   Boundary("End 1895");
  };
  Sequence("Year of death 1896")
  {
   Boundary("Start 1896");
   Phase("1896 walrus")
   {
    R_Date("TRa-25064",535,13);
    R_Date("TRa-25069",582,13);
    R_Date("TRa-25065",599,15);
    R_Date("TRa-25068",603,14);
    R_Date("TRa-25063",569,21);
   };
```

```
  C_Date("1896",1896,1);
  Boundary("End 1896");
 };
 Sequence("Year of death 1926")
 {
  Boundary("Start 1926");
  Phase("1926 walrus")
  {
   R_Date("TRa-25060",553,16);
   R_Date("TRa-25061",563,13);
   R_Date("TRa-25062",565,18);
  };
  C_Date("1926",1926,1);
  Boundary("End 1926");
 };
};
```

**2.1.2 Baffin Bay, including date of death and estimated date of birth**

```
Plot()
{
 Curve("Marine20", "marine20.14c");
 Delta_R("BAF",U(-500,500));
 Sequence("Year of death 1853")
 {
  Boundary("Start 1853");
  C_Date("1853 adult",1828,9);
  R_Date("TRa-22945",649,12);
  C_Date("1853",1853,2);
  Boundary("End 1853");
 };
 Sequence("Year of death 1854")
 {
  Boundary("Start 1854");
  C_Date("1854 adults",1829,9);
  Phase("1854 walrus")
  {
   R_Date("TRa-22943",725,15);
   R_Date("TRa-22944",737,13);
  };
  C_Date("1854",1854,2);
  Boundary("End 1854");
 };
 Sequence("Year of death 1895")
 {
```

```
 Boundary("Start 1895");
 C_Date("1895 adult",1870,8);
 R_Date("TRa-25066",582,13);
 C_Date("1895",1895,1);
 Boundary("End 1895");
};
Sequence("Year of death 1896")
{
 Boundary("Start 1896");
 C_Date("1896 adults",1871,8);
 Phase("1896 walrus")
 {
  R_Date("TRa-25064",535,13);
  R_Date("TRa-25069",582,13);
  R_Date("TRa-25065",599,15);
  R_Date("TRa-25068",603,14);
  R_Date("TRa-25063",569,21);
 };
 C_Date("1896",1896,1);
 Boundary("End 1896");
};
Sequence("Year of death 1926 adults")
{
 Boundary("Start 1926 adults");
 C_Date("1926 adults DOB",1901,8);
 Phase("1926 walrus")
 {
  R_Date("TRa-25060",553,16);
  R_Date("TRa-25062",565,18);
 };
 C_Date("1926 adults DOD",1926,1);
 Boundary("End 1926 adults");
};
Sequence("Year of death 1926 juvenile")
{
 Boundary("Start 1926 juvenile");
 C_Date("1926 juvenile DOB",1919,2);
 R_Date("TRa-25061",563,13);
 C_Date("1926 juvenile DOD",1926,1);
 Boundary("End 1926 juvenile");
};
};
```

**2.1.3 Baffin Bay, including date of death and estimated date of birth; pre-1950 dates from this study and Dyke et al. 2019**

```
Plot()
 {
  Curve("Marine20", "marine20.14c");
  Delta_R("BAF",U(-500,500));
  Sequence("Year of death 1853")
  {
   Boundary("Start 1853");
   C_Date("1853 adult",1828,9);
   R_Date("TRa-22945",649,12);
   C_Date("1853",1853,2);
   Boundary("End 1853");
  };
  Sequence("Year of death 1854")
  {
   Boundary("Start 1854");
   C_Date("1854 adults",1829,9);
   Phase("1854 walrus")
   {
    R_Date("TRa-22943",725,15);
    R_Date("TRa-22944",737,13);
   };
   C_Date("1854",1854,2);
   Boundary("End 1854");
  };
  Sequence("Year of death 1895")
  {
   Boundary("Start 1895");
   C_Date("1895 adult",1870,8);
   R_Date("TRa-25066",582,13);
   C_Date("1895",1895,1);
   Boundary("End 1895");
  };
  Sequence("Year of death 1896")
  {
   Boundary("Start 1896");
   C_Date("1896 adults",1871,8);
   Phase("1896 walrus")
   {
    R_Date("TRa-25064",535,13);
    R_Date("TRa-25069",582,13);
    R_Date("TRa-25065",599,15);
    R_Date("TRa-25068",603,14);
```

```
   R_Date("TRa-25063",569,21);
  };
  C_Date("1896",1896,1);
  Boundary("End 1896");
 };
Sequence("Year of death 1924 ontology uncertain")
 {
  Boundary("Start 1924 ontology uncertain");
  C_Date("1924 ontology uncertain DOB",1904,10);
  R_Date("UCIAMS-168854",755,15);
  C_Date("1924 ontology uncertain DOD",1924,1);
  Boundary("End 1924 ontology uncertain");
 };
 Sequence("Year of death 1926 adults")
 {
  Boundary("Start 1926 adults");
  C_Date("1926 adults DOB",1901,8);
  Phase("1926 walrus")
  {
   R_Date("TRa-25060",553,16);
   R_Date("TRa-25062",565,18);
  };
  C_Date("1926 adults DOD",1926,1);
  Boundary("End 1926 adults");
 };
 Sequence("Year of death 1926 juvenile")
 {
  Boundary("Start 1926 juvenile");
  C_Date("1926 juvenile DOB",1919,2);
  R_Date("TRa-25061",563,13);
  C_Date("1926 juvenile DOD",1926,1);
  Boundary("End 1926 juvenile");
 };
};
```

**2.2.1 West Greenland, including date of death only**

```
Plot()
 {
  Curve("Marine20", "marine20.14c");
  Delta_R("WGL",U(-500,500));
  Sequence("Year of death 1898")
  {
   Boundary("Start 1898");
   R_Date("TRa-25591",604,13);
```

```
  C_Date("1898",1898,1);
  Boundary("End 1898");
 };
 Sequence("Year of death 1895")
 {
  Boundary("Start 1895");
  R_Date("TRa-25067",611,20);
  C_Date("1895",1895,1);
  Boundary("End 1895");
 };
};
```

**2.2.2 West Greenland, including date of death and estimated date of birth**

```
Plot()
{
 Curve("Marine20", "marine20.14c");
 Delta_R("WGL",U(-500,500));
 Sequence("Year of death 1898")
 {
  Boundary("Start 1898");
  C_Date("1898 adult DOB",1873,8);
  R_Date("TRa-25591",604,13);
  C_Date("1898 DOD",1898,1);
  Boundary("End 1898");
 };
 Sequence("Year of death 1895")
 {
  Boundary("Start 1895");
  C_Date("1895 juvenile DOB",1888,2);
  R_Date("TRa-25067",611,20);
  C_Date("1895 DOD",1895,1);
  Boundary("End 1895");
 };
};
```

**2.3.1 Foxe Basin, including date of death only**

```
Plot()
{
 Curve("Marine20", "marine20.14c");
 Delta_R("SFO",U(-500,500));
 Sequence("Year of death 1922")
 {
  Boundary("Start 1922");
  Phase("phase 1922")
```

```
  {
   R_Date("TRa-25596",687,16);
   R_Date("TRa-25595",646,18);
  };
  C_Date("1922",1922,1);
  Boundary("End 1922");
 };
};
```

**2.3.2 Foxe Basin, including date of death and estimated date of birth**

```
Plot()
 {
  Curve("Marine20", "marine20.14c");
  Delta_R("SFO",U(-500,500));
  Sequence("Year of death 1922")
  {
   Boundary("Start 1922");
   C_Date("1922 juveniles DOB",1915,2);
   Phase("phase 1922")
   {
    R_Date("TRa-25596",687,16);
    R_Date("TRa-25595",646,18);
   };
   C_Date("1922 juveniles DOD",1922,1);
   Boundary("End 1922");
  };
 };
```

**2.3.3 Foxe Basin, including date of death and estimated date of birth; pre-1950 dates from this study and Dyke et al. 2019**

```
Plot()
 {
  Curve("Marine20", "marine20.14c");
  Delta_R("SFO",U(-500,500));
  Sequence("Year of death 1922")
  {
   Boundary("Start 1922");
   C_Date("1922 juveniles DOB",1915,2);
   Phase("phase 1922")
   {
    R_Date("TRa-25596",687,16);
    R_Date("TRa-25595",646,18);
   };
   C_Date("1922 juveniles DOD",1922,1);
```

```
  Boundary("End 1922");
 };
 Sequence("Year of death 1949")
 {
  Boundary("Start 1949");
  C_Date("1949 ontology unknown DOB",1929,10);
  Phase("phase 1949")
  {
   R_Date("UCIAMS-168835",595,15);
   R_Date("UCIAMS-168834",655,15);
   R_Date("UCIAMS-168833",685,15);
  };
  C_Date("1949 ontology unknown DOD",1949,1);
  Boundary("End 1949");
 };
};
```

**2.4.1 Hudson Strait/Hudson Bay, including date of death only**

```
Plot()
{
 Curve("Marine20", "marine20.14c");
 Delta_R("HUD",U(-500,500));
 Sequence("Year of death 1885")
 {
  Boundary("Start 1885");
  Phase("1885 walrus")
  {
   R_Date("TRa-24149",671,20);
   R_Date("TRa-24150",639,13);
  };
  C_Date("1885",1885,2);
  Boundary("End 1885");
 };
 Sequence("Year of death 1886")
 {
  Boundary("Start 1886");
  R_Date("TRa-24148",609,12);
  C_Date("1886",1886,1);
  Boundary("End 1886");
 };
 Sequence("Year of death 1910")
 {
  Boundary("Start 1910");
  R_Date("TRa-24151",615,12);
```

```
  C_Date("1910",1910,1);
  Boundary("End 1910");
 };
 };
```

**2.4.2 Hudson Strait/Hudson Bay, including date of death and estimated date of birth**

```
Plot()
 {
  Curve("Marine20", "marine20.14c");
  Delta_R("HUD",U(-500,500));
  Sequence("Year of death 1885")
  {
   Boundary("Start 1885");
   C_Date("1885 juveniles DOB",1878,3);
   Phase("1885 walrus")
   {
    R_Date("TRa-24149",671,20);
    R_Date("TRa-24150",639,13);
   };
   C_Date("1885 DOD",1885,2);
   Boundary("End 1885");
  };
  Sequence("Year of death 1886")
  {
   Boundary("Start 1886");
   C_Date("1886 adult DOB",1861,8);
   R_Date("TRa-24148",609,12);
   C_Date("1886 DOD",1886,1);
   Boundary("End 1886");
  };
  Sequence("Year of death 1910")
  {
   Boundary("Start 1910");
   C_Date("1910 adult DOB",1885,8);
   R_Date("TRa-24151",615,12);
   C_Date("1910 DOD",1910,1);
   Boundary("End 1910");
 };
 };
```

**2.4.3 Hudson Strait/Hudson Bay, including date of death and estimated date of birth; pre-1950 dates from this study and Dyke et al. 2019**

```
Plot()
```

```
{
 Curve("Marine20", "marine20.14c");
 Delta_R("HUD",U(-500,500));
 Sequence("Year of death 1885 ontology uncertain")
{
 Boundary("Start 1885 ontology uncertain");
 C_Date("1885 ontology uncertain DOB",1865,10);
 R_Date("UCIAMS-168836",670,15);
 C_Date("1885 ontology uncertain DOD",1885,1);
 Boundary("End 1885 ontology uncertain");
 };
Sequence("Year of death 1885 juveniles")
 {
  Boundary("Start 1885 juveniles");
  C_Date("1885 juveniles DOB",1878,3);
  Phase("1885 juvenile walruses")
  {
   R_Date("TRa-24149",671,20);
   R_Combine("CMH36")
  {
   R_Date("CMH36_TRa-24150",639,13);
   R_Date("CMH36_UCIAMS-168850",665,15);
  };
  };
  C_Date("1885 juveniles DOD",1885,2);
  Boundary("End 1885 juveniles");
 };
  Sequence("Year of death 1886 adult")
 {
  Boundary("Start 1886 adult");
  C_Date("1886 adult DOB",1861,8);
  R_Date("TRa-24148",609,12);
  C_Date("1886 adult DOD",1886,1);
  Boundary("End 1886 adult");
 };
Sequence("Year of death 1886 ontology uncertain")
 {
  Boundary("Start 1886 ontology uncertain");
  C_Date("1886 ontology uncertain DOB",1866,10);
  R_Date("UCIAMS-168857",630,15);
  C_Date("1886 ontology uncertain DOD",1886,1);
  Boundary("End 1886 ontology uncertain");
 };
 Sequence("Year of death 1910")
```

```
 {
  Boundary("Start 1910");
  C_Date("1910 adult DOB",1885,8);
  R_Date("TRa-24151",615,12);
  C_Date("1910 DOD",1910,1);
  Boundary("End 1910");
 };
Sequence("Year of death 1923")
 {
  Boundary("Start 1923 ontology uncertain");
  C_Date("1923 ontology uncertain DOB",1903,10);
  R_Date("UCIAMS-168851",595,15);
  C_Date("1923 ontology uncertain DOD",1923,1);
  Boundary("End 1923 ontology uncertain");
 };
Sequence("Year of death 1924")
 {
  Boundary("Start 1924 ontology uncertain");
  C_Date("1924 ontology uncertain DOB",1904,10);
  Phase("phase 1924 ontology uncertain")
  {
   R_Date("UCIAMS-168852",660,15);
   R_Date("UCIAMS-168855",670,15);
   R_Date("UCIAMS-168853",690,15);
   R_Date("UCIAMS-168856",695,15);
  };
  C_Date("1924 ontology uncertain DOD",1924,1);
  Boundary("End 1924 ontology uncertain");
 };
Sequence("Year of death 1928")
 {
  Boundary("Start 1928 ontology uncertain");
  C_Date("1928 ontology uncertain DOB",1908,10);
  Phase("phase 1928 ontology uncertain")
  {
   R_Date("UCIAMS-185717",625,15);
   R_Date("UCIAMS-168831",705,15);
  };
  C_Date("1928 ontology uncertain DOD",1928,1);
  Boundary("End 1928 ontology uncertain");
 };
Sequence("Year of death 1945")
 {
  Boundary("Start 1945 ontology uncertain");
```

```
  C_Date("1945 ontology uncertain DOB",1925,10);
  R_Date("UCIAMS-168832",635,15);
  C_Date("1945 ontology uncertain DOD",1945,1);
  Boundary("End 1945 ontology uncertain");
 };
};
```

**2.5.1 Pooled Greenland (excluding East Greenland) and Canadian Arctic, including date of death only**

```
Plot()
 {
  Curve("Marine20", "marine20.14c");
  Delta_R("Greenl_Can_Arct",U(-500,500));
  Sequence("Baffin Year of death 1853")
  {
   Boundary("Start BAF 1853");
   R_Date("TRa-22945",649,12);
   C_Date("1853",1853,2);
   Boundary("End BAF 1853");
  };
  Sequence("Baffin Year of death 1854")
  {
   Boundary("Start BAF 1854");
   Phase("BAF 1854 walrus")
   {
    R_Date("TRa-22943",725,15);
    R_Date("TRa-22944",737,13);
   };
   C_Date("1854",1854,2);
   Boundary("End BAF 1854");
  };
  Sequence("Hudson Strait Year of death 1885")
  {
   Boundary("Start HUD 1885");
   Phase("HUD 1885 walrus")
   {
    R_Date("TRa-24149",671,20);
    R_Date("TRa-24150",639,13);
   };
   C_Date("1885",1885,2);
   Boundary("End HUD 1885");
  };
  Sequence("Hudson Strait Year of death 1886")
  {
```

```
  Boundary("Start HUD 1886");
  R_Date("TRa-24148",609,12);
  C_Date("1886",1886,1);
  Boundary("End HUD 1886");
 };
 Sequence("Baffin Year of death 1895")
 {
  Boundary("Start BAF 1895");
  R_Date("TRa-25066",582,13);
  C_Date("Baffin 1895",1895,1);
  Boundary("End BAF 1895");
 };
Sequence("Western Greenland Year of death 1895")
 {
  Boundary("Start WG 1895");
  R_Date("TRa-25067",611,20);
  C_Date("1895",1895,1);
  Boundary("End WG 1895");
 };
 Sequence("Baffin Year of death 1896")
 {
  Boundary("Start BAF 1896");
  Phase("BAF 1896 walrus")
  {
   R_Date("TRa-25064",535,13);
   R_Date("TRa-25069",582,13);
   R_Date("TRa-25065",599,15);
   R_Date("TRa-25068",603,14);
   R_Date("TRa-25063",569,21);
  };
  C_Date("1896",1896,1);
  Boundary("End BAF 1896");
 };
 Sequence("Western Greenland Year of death 1898")
 {
  Boundary("Start WG 1898");
  R_Date("TRa-25591",604,13);
  C_Date("1898",1898,1);
  Boundary("End WG 1898");
 };
Sequence("Hudson Strait_Hudson Bay Year of death 1910")
 {
  Boundary("Start HUD 1910");
  R_Date("TRa-24151",615,12);
```

```
 C_Date("1910",1910,1);
 Boundary("End HUD 1910");
};
Sequence("Foxe Basin Year of death 1922")
{
 Boundary("Start FOX 1922");
 Phase("FOX 1922 walrus")
 {
  R_Date("TRa-25596",687,16);
  R_Date("TRa-25595",646,18);
 };
 C_Date("Foxe 1922",1922,1);
 Boundary("End FOX 1922");
};
Sequence("Baffin Year of death 1926")
{
 Boundary("Start BAF 1926");
 Phase("BAF 1926 walrus")
 {
  R_Date("TRa-25060",553,16);
  R_Date("TRa-25061",563,13);
  R_Date("TRa-25062",565,18);
 };
 C_Date("1926",1926,1);
 Boundary("End BAF 1926");
};
};
```

**2.5.2 Pooled Greenland (excluding East Greenland) and Canadian Arctic, including date of death and estimated date of birth**

```
Plot()
{
 Curve("Marine20", "marine20.14c");
 Delta_R("BAF",U(-500,500));
 Sequence("BAF Year of death 1853")
 {
  Boundary("Start BAF 1853");
  C_Date("BAF 1853 adult DOB",1828,9);
  R_Date("TRa-22945",649,12);
  C_Date("BAF 1853 adult DOD",1853,2);
  Boundary("End BAF 1853");
 };
 Sequence("BAF Year of death 1854")
```

```
{
 Boundary("Start BAF 1854");
 C_Date("BAF 1854 adults DOB",1829,9);
 Phase("phase BAF 1854 walrus")
 {
  R_Date("TRa-22943",725,15);
  R_Date("TRa-22944",737,13);
 };
 C_Date("BAF 1854 adults DOD",1854,2);
 Boundary("End BAF 1854");
};
Sequence("HUD Year of death 1885")
{
 Boundary("Start HUD 1885");
 C_Date("HUD 1885 juveniles DOB",1878,3);
 Phase("phase HUD 1885 walrus")
 {
  R_Date("TRa-24149",671,20);
  R_Date("TRa-24150",639,13);
 };
 C_Date("HUD 1885 juveniles DOD",1885,2);
 Boundary("End HUD 1885");
};
Sequence("HUD Year of death 1886")
{
 Boundary("Start HUD 1886");
 C_Date("HUD 1886 adult DOB",1861,8);
 R_Date("TRa-24148",609,12);
 C_Date("HUD 1886 adult DOD",1886,1);
 Boundary("End HUD 1886");
};
Sequence("BAF Year of death 1895 adult")
{
 Boundary("Start BAF 1895");
 C_Date("BAF 1895 adult DOB",1870,8);
 R_Date("TRa-25066",582,13);
 C_Date("BAF 1895 adult DOD",1895,1);
 Boundary("End BAF 1895");
};
Sequence("WGL Year of death 1895 juvenile")
{
 Boundary("Start WGL 1895");
 C_Date("WGL 1895 juvenile DOB",1888,2);
 R_Date("TRa-25067",611,20);
```

```
 C_Date("WGL 1895 juvenile DOD",1895,1);
 Boundary("End WGL 1895");
};
Sequence("BAF Year of death 1896")
{
 Boundary("Start BAF 1896");
 C_Date("BAF 1896 adults DOB",1871,8);
 Phase("phase BAF 1896 walrus")
 {
  R_Date("TRa-25064",535,13);
  R_Date("TRa-25069",582,13);
  R_Date("TRa-25065",599,15);
  R_Date("TRa-25068",603,14);
  R_Date("TRa-25063",569,21);
 };
 C_Date("BAF 1896 adults DOD",1896,1);
 Boundary("End BAF 1896");
};
Sequence("WGL Year of death 1898")
{
 Boundary("Start WGL 1898");
 C_Date("WGL 1898 adult DOB",1873,8);
 R_Date("TRa-25591",604,13);
 C_Date("WGL 1898 DOD",1898,1);
 Boundary("End WGL 1898");
};
Sequence("HUD Year of death 1910")
{
 Boundary("Start HUD 1910");
 C_Date("HUD 1910 adult DOB",1885,8);
 R_Date("TRa-24151",615,12);
 C_Date("HUD 1910 adult DOD",1910,1);
 Boundary("End HUD 1910");
};
Sequence("FOX Year of death 1922")
{
 Boundary("Start FOX 1922");
 C_Date("FOX 1922 juveniles DOB",1915,2);
 Phase("phase FOX 1922")
 {
  R_Date("TRa-25596",687,16);
  R_Date("TRa-25595",646,18);
 };
 C_Date("FOX 1922 juveniles DOD",1922,1);
```

```
  Boundary("End FOX 1922");
 };
 Sequence("BAF Year of death 1926 adults")
 {
  Boundary("Start BAF 1926 adults");
  C_Date("BAF 1926 adults DOB",1901,8);
  Phase("phase BAF 1926 walrus")
  {
   R_Date("TRa-25060",553,16);
   R_Date("TRa-25062",565,18);
  };
  C_Date("BAF 1926 adults DOD",1926,1);
  Boundary("End BAF 1926 adults");
 };
 Sequence("BAF Year of death 1926 juvenile")
 {
  Boundary("Start BAF 1926 juvenile");
  C_Date("BAF 1926 juvenile DOB",1919,2);
  R_Date("TRa-25061",563,13);
  C_Date("BAF 1926 juvenile DOD",1926,1);
  Boundary("End BAF 1926 juvenile");
 };
};
```

**2.5.3 Pooled Greenland (excluding East Greenland) and Canadian Arctic, including date of death and estimated date of birth; pre-1950 dates from this study and Dyke et al. 2019**

```
Plot()
 {
  Curve("Marine20", "marine20.14c");
  Delta_R("Greenl_Can_Arct",U(-500,500));
Sequence("BAF Year of death 1853")
  {
   Boundary("Start BAF 1853");
   C_Date("BAF 1853 adult DOB",1828,9);
   R_Date("TRa-22945",649,12);
   C_Date("BAF 1853 adult DOD",1853,2);
   Boundary("End BAF 1853");
  };
Sequence("BAF Year of death 1854")
  {
   Boundary("Start BAF 1854");
   C_Date("BAF 1854 adults DOB",1829,9);
   Phase("phase BAF 1854 walrus")
   {
```

```
    R_Date("TRa-22943",725,15);
    R_Date("TRa-22944",737,13);
   };
   C_Date("BAF 1854 adults DOD",1854,2);
   Boundary("End BAF 1854");
  };
Sequence("HUD Year of death 1885 ontology uncertain")
 {
  Boundary("Start HUD 1885 ontology uncertain");
  C_Date("HUD 1885 ontology uncertain DOB",1865,10);
  R_Date("UCIAMS-168836",670,15);
  C_Date("HUD 1885 ontology uncertain DOD",1885,1);
  Boundary("End HUD 1885 ontology uncertain");
  };
Sequence("HUD Year of death 1885 juveniles")
  {
   Boundary("Start HUD 1885 juveniles");
   C_Date("HUD 1885 juveniles DOB",1878,3);
   Phase("phase HUD 1885 juvenile walruses")
   {
    R_Date("TRa-24149",671,20);
    R_Combine("CMH36")
   {
    R_Date("CMH36_TRa-24150",639,13);
    R_Date("CMH36_UCIAMS-168850",665,15);
   };
   };
   C_Date("HUD 1885 juveniles DOD",1885,2);
   Boundary("End HUD 1885 juveniles");
  };
Sequence("HUD Year of death 1886 adult")
  {
   Boundary("Start HUD 1886 adult");
   C_Date("HUD 1886 adult DOB",1861,8);
   R_Date("TRa-24148",609,12);
   C_Date("HUD 1886 adult DOD",1886,1);
   Boundary("End HUD 1886 adult");
  };
Sequence("HUD Year of death 1886 ontology uncertain")
  {
   Boundary("Start HUD 1886 ontology uncertain");
   C_Date("HUD 1886 ontology uncertain DOB",1866,10);
   R_Date("UCIAMS-168857",630,15);
   C_Date("HUD 1886 ontology uncertain DOD",1886,1);
```

```
   Boundary("End HUD 1886 ontology uncertain");
  };
Sequence("BAF Year of death 1895 adult")
  {
   Boundary("Start BAF 1895");
   C_Date("BAF 1895 adult DOB",1870,8);
   R_Date("TRa-25066",582,13);
   C_Date("BAF 1895 adult DOD",1895,1);
   Boundary("End BAF 1895");
  };
Sequence("WGL Year of death 1895 juvenile")
  {
   Boundary("Start WGL 1895");
   C_Date("WGL 1895 juvenile DOB",1888,2);
   R_Date("TRa-25067",611,20);
   C_Date("WGL 1895 juvenile DOD",1895,1);
   Boundary("End WGL 1895");
  };
Sequence("BAF Year of death 1896")
  {
   Boundary("Start BAF 1896");
   C_Date("BAF 1896 adults DOB",1871,8);
   Phase("phase BAF 1896 walrus")
   {
    R_Date("TRa-25064",535,13);
    R_Date("TRa-25069",582,13);
    R_Date("TRa-25065",599,15);
    R_Date("TRa-25068",603,14);
    R_Date("TRa-25063",569,21);
   };
   C_Date("BAF 1896 adults DOD",1896,1);
   Boundary("End BAF 1896");
  };
Sequence("WGL Year of death 1898")
  {
   Boundary("Start WGL 1898");
   C_Date("WGL 1898 adult DOB",1873,8);
   R_Date("TRa-25591",604,13);
   C_Date("WGL 1898 DOD",1898,1);
   Boundary("End WGL 1898");
  };
Sequence("HUD Year of death 1910")
  {
   Boundary("Start HUD 1910");
```

```
   C_Date("HUD 1910 adult DOB",1885,8);
   R_Date("TRa-24151",615,12);
   C_Date("HUD 1910 adult DOD",1910,1);
   Boundary("End HUD 1910");
  };
Sequence("FOX Year of death 1922")
  {
   Boundary("Start FOX 1922");
   C_Date("FOX 1922 juveniles DOB",1915,2);
   Phase("phase FOX 1922")
   {
    R_Date("TRa-25596",687,16);
    R_Date("TRa-25595",646,18);
   };
   C_Date("FOX 1922 juveniles DOD",1922,1);
   Boundary("End FOX 1922");
  };
Sequence("HUD Year of death 1923")
  {
   Boundary("Start HUD 1923 ontology uncertain");
   C_Date("HUD 1923 ontology uncertain DOB",1903,10);
   R_Date("UCIAMS-168851",595,15);
   C_Date("HUD 1923 ontology uncertain DOD",1923,1);
   Boundary("End HUD 1923 ontology uncertain");
  };
Sequence("BAF Year of death 1924 ontology uncertain")
  {
   Boundary("Start BAF 1924 ontology uncertain");
   C_Date("BAF 1924 ontology uncertain DOB",1904,10);
   R_Date("UCIAMS-168854",755,15);
   C_Date("BAF 1924 ontology uncertain DOD",1924,1);
   Boundary("End BAF 1924 ontology uncertain");
  };
Sequence("HUD Year of death 1924 ontology uncertain")
  {
   Boundary("Start HUD 1924 ontology uncertain");
   C_Date("HUD 1924 ontology uncertain DOB",1904,10);
   Phase("phase HUD 1924 ontology uncertain")
   {
    R_Date("UCIAMS-168852",660,15);
    R_Date("UCIAMS-168855",670,15);
    R_Date("UCIAMS-168853",690,15);
    R_Date("UCIAMS-168856",695,15);
   };
```

```
  C_Date("HUD 1924 ontology uncertain DOD",1924,1);
  Boundary("End HUD 1924 ontology uncertain");
 };
Sequence("BAF Year of death 1926 adults")
 {
  Boundary("Start BAF 1926 adults");
  C_Date("BAF 1926 adults DOB",1901,8);
  Phase("phase BAF 1926 walrus")
  {
   R_Date("TRa-25060",553,16);
   R_Date("TRa-25062",565,18);
  };
  C_Date("BAF 1926 adults DOD",1926,1);
  Boundary("End BAF 1926 adults");
 };
Sequence("BAF Year of death 1926 juvenile")
 {
  Boundary("Start BAF 1926 juvenile");
  C_Date("BAF 1926 juvenile DOB",1919,2);
  R_Date("TRa-25061",563,13);
  C_Date("BAF 1926 juvenile DOD",1926,1);
  Boundary("End BAF 1926 juvenile");
 };
Sequence("HUD Year of death 1928")
 {
  Boundary("StartHUD 1928 ontology uncertain");
  C_Date("HUD 1928 ontology uncertain DOB",1908,10);
  Phase("phase HUD 1928 ontology uncertain")
  {
   R_Date("UCIAMS-185717",625,15);
   R_Date("UCIAMS-168831",705,15);
  };
  C_Date("HUD 1928 ontology uncertain DOD",1928,1);
  Boundary("End HUD 1928 ontology uncertain");
 };
Sequence("HUD Year of death 1945")
 {
  Boundary("Start HUD 1945 ontology uncertain");
  C_Date("HUD 1945 ontology uncertain DOB",1925,10);
  R_Date("UCIAMS-168832",635,15);
  C_Date("HUD 1945 ontology uncertain DOD",1945,1);
  Boundary("End HUD 1945 ontology uncertain");
 };
Sequence("FOX Year of death 1949")
```

```
 {
  Boundary("Start FOX 1949");
  C_Date("FOX 1949 ontology unknown DOB",1929,10);
  Phase("phase FOX 1949")
  {
   R_Date("UCIAMS-168835",595,15);
   R_Date("UCIAMS-168834",655,15);
   R_Date("UCIAMS-168833",685,15);
  };
  C_Date("FOX 1949 ontology unknown DOD",1949,1);
  Boundary("End FOX 1949");
 };
};
```

**2.6.1 East Greenland, including date of death only**

```
Plot()
 {
  Curve("Marine20", "marine20.14c");
  Delta_R("EGL",U(-500,500));
  Sequence("Year of death 1924")
  {
   Boundary("Start 1924");
   R_Date("TRa-25593",569,13);
   C_Date("1924",1924,1);
   Boundary("End 1924");
  };
 Sequence("Year of death 1927")
  {
   Boundary("Start 1927");
   R_Date("TRa-25587",605,23);
   C_Date("1927",1927,1);
   Boundary("End 1927");
  };
 Sequence("Year of death 1929")
  {
   Boundary("Start 1929");
   Phase("1929 walruses")
   {
    R_Date("TRa-25585",561,16);
    R_Date("TRa-25588",558,16);
    R_Date("TRa-25586",567,15);
```

```
  };
  C_Date("1929 DOD",1929,3);
  Boundary("End 1929");
 };
 Sequence("Year of death 1909")
 {
  Boundary("Start 1909");
  R_Date("TRa-25589",631,21);
  C_Date("1909",1909,4);
  Boundary("End 1909");
 };
 };
```

**2.6.2 East Greenland, including date of death and estimated date of birth**

```
Plot()
{
 Curve("Marine20", "marine20.14c");
 Delta_R("EGL",U(-500,500));
Sequence("Year of death 1924")
 {
  Boundary("Start 1924");
  C_Date("1924 adult DOB",1899,8);
  R_Date("TRa-25593",569,13);
  C_Date("1924 DOD",1924,1);
  Boundary("End 1924");
 };
 Sequence("Year of death 1927")
 {
  Boundary("Start 1927");
  C_Date("1927 adult DOB",1902,8);
  R_Date("TRa-25587",605,23);
  C_Date("1927 DOD",1927,1);
  Boundary("End 1927");
 };
 Sequence("Year of death 1929 adults")
 {
  Boundary("Start 1929 adults");
  C_Date("1929 adults DOB",1904,10);
  Phase("1929 adults")
  {
   R_Date("TRa-25585",561,16);
   R_Date("TRa-25588",558,16);
  };
  C_Date("1929 adults DOD",1929,3);
```

```
 Boundary("End 1929 adults");
};
Sequence("Year of death 1929 juvenile")
{
 Boundary("Start 1929 juvenile");
 C_Date("1929 juvenile DOB",1922,2);
 R_Date("TRa-25586",567,15);
 C_Date("1929 juvenile DOD",1929,1);
 Boundary("End 1929 juvenile");
};
Sequence("Year of death 1909")
{
 Boundary("Start 1909");
 C_Date("1909 adult DOB",1884,11);
 R_Date("TRa-25589",631,21);
 C_Date("1909 DOD",1909,4);
 Boundary("End 1909");
};
};
```

**2.7.1 Iceland, including date of death only**

```
Plot()
 {
  Curve("Marine20", "marine20.14c");
  Delta_R("ICE",U(-500,500));
  Sequence("Year of death 1900")
  {
   Boundary("Start 1900");
   R_Date("TRa-25590",576,14);
   C_Date("1900",1900,1);
   Boundary("End 1900");
  };
 };
```

**2.7.2 Iceland, including date of death and estimated date of birth**

```
Plot()
 {
  Curve("Marine20", "marine20.14c");
  Delta_R("ICE",U(-500,500));
  Sequence("Year of death 1900")
  {
   Boundary("Start 1900");
   C_Date("1900 adult DOB",1875,8);
   R_Date("TRa-25590",576,14);
```

```
  C_Date("1900 DOD",1900,1);
  Boundary("End 1900");
 };
};
```

**2.8.1 Franz Josef Land, including date of death only**

```
Plot()
{
 Curve("Marine20", "marine20.14c");
 Delta_R("FJL",U(-500,500));
 Sequence("Year of death 1894 to 1897")
 {
  Boundary("Start 1897");
  Phase("phase 1897")
  {
   R_Date("TRa-22952",472,15);
   R_Date("TRa-22947",494,15);
  };
  C_Date("1897",1897,4);
  Boundary("End 1897");
 };
};
```

**2.8.2 Franz Josef Land, including date of death and estimated date of birth**

```
Plot()
{
 Curve("Marine20", "marine20.14c");
 Delta_R("FJL",U(-500,500));
 Sequence("Year of death 1894 to 1897 adult")
 {
  Boundary("Start 1897 adult");
  C_Date("1897 adult DOB",1872,11);
  R_Date("TRa-22947",494,15);
  C_Date("1897 adult DOD",1897,4);
  Boundary("End 1897 adult");
 };
Sequence("Year of death 1894 to 1897 neonatal")
 {
  Boundary("Start 1897 neonatal");
  C_Date("1897 neonatal DOB",1895,4);
  R_Date("TRa-22952",472,15);
  C_Date("1897 neonatal DOD",1897,4);
  Boundary("End 1897 neonatal");
 };
```

```
};
```

**2.9.1 Pacific, including date of death only**

```
Plot()
 {
  Curve("Marine20", "marine20.14c");
  Delta_R("PAC",U(-500,500));
 Sequence("Year of death 1865")
  {
   Boundary("Start 1865");
   R_Date("TRa-22951",843,13);
   C_Date("1865",1865,1);
   Boundary("End 1865");
 };
 Sequence("Year of death 1897")
  {
   Boundary("Start 1897");
   R_Date("TRa-22950",1026,16);
   C_Date("1897",1897,2);
   Boundary("End 1897");
  };
 };
```

**2.9.2 Pacific, including date of death and estimated date of birth**

```
Plot()
 {
  Curve("Marine20", "marine20.14c");
  Delta_R("PAC",U(-500,500));
 Sequence("Year of death 1865")
  {
   Boundary("Start 1865");
   C_Date("1865 juvenile DOB",1858,2);
   R_Date("TRa-22951",843,13);
   C_Date("1865 DOD",1865,1);
   Boundary("End 1865");
 };
  Sequence("Year of death 1897")
  {
   Boundary("Start 1897");
   C_Date("1897 adult DOB",1872,9);
   R_Date("TRa-22950",1026,16);
   C_Date("1897 DOD",1897,2);
   Boundary("End 1897");
  };
```

```
};
```

**2.10.1 Calibration using ΔR of 17±12 with no constraints based on archaeological dating**

```
Plot()
{
 Curve("Marine20", "marine20.14c");
 Delta_R("Pooled", 17,12);
 R_Date("TRa-25177",1229,13);
 R_Date("TRa-25175",1389,21);
 R_Date("TRa-25176",1388,14);
 R_Date("TRa-25178",1363,20);
 R_Date("TRa-25883",1226,13);
 R_Date("TRa-25884",1321,12);
 R_Date("TRa-25886",1364,34);
 R_Date("TRa-25887",1255,11);
 R_Date("TRa-25885",1351,12);
 R_Date("TRa-25179",1312,13);
 R_Date("TRa-25888",1313,14);
 R_Date("TRa-25180",1399,16);
};
```

**2.10.2 Calibration including archaeological dating**

This code includes both options mentioned in the text, 1) using the ΔR of 17±12 calculated for historical specimens from Greenland (excluding Eastern Greenland) and the Canadian Arctic, together with archaeological dating of the archaeological find contexts and 2) allowing OxCal to estimate a ΔR based on the nominal dating of the archaeological find contexts (depending on which Delta_R command is commented out).

```
Plot()
{
 Curve("Marine20", "marine20.14c");
 //Delta_R("Modern DeltaR W Greenland and Canadian Arctic", 17,12);
 Delta_R("DeltaR old walrus",U(-400,400));
 Sequence("Trondheim 1111")
 {
  Boundary("Start Trondheim 1111");
  R_Date("TRa-25180", 1399, 16);
  C_Date("1111", 1111, 1);
  Boundary("End Trondheim 1111");
 };
 Label("Trondheim post 1600:");
 R_Date("TRa-25175", 1389, 21);
 Sequence("Trondheim 1225-1275")
 {
```

```
  Boundary("Start Trondheim 1225-1275");
  Phase("Phase Trondheim 1225-1275")
  {
          R_Date("TRa-25176", 1388, 14);
          R_Date("TRa-25883", 1226, 13);
  };
  C_Date("1225-1275", 1250, 13);
  Boundary("End Trondheim 1225-1275");
 };
 Sequence("Trondheim 1100-1150")
 {
  Boundary("Start Trondheim 1100-1150");
  R_Date("TRa-25886", 1364, 34);
  C_Date("1100-1150", 1125, 13);
  Boundary("End Trondheim 1100-1150");
 };
 Sequence("Trondheim 1150-1175")
 {
  Boundary("Start Trondheim 1150-1175");
  R_Date("TRa-25178", 1363, 20);
  C_Date("1150-1175", 1163, 6);
  Boundary("End Trondheim 1150-1175");
 };
 R_Date("TRa-25885", 1351, 12);
 Sequence("Trondheim 1175-1225")
 {
  Boundary("Start Trondheim 1175-1225");
  R_Date("TRa-25884", 1321, 12);
  C_Date("1175-1225", 1200, 13);
  Boundary("End Trondheim 1175-1225");
 };
 R_Date("TRa-25888", 1313, 14);
 Sequence("Trondheim 12th century")
 {
  Boundary("Start Trondheim 12th century");
  R_Date("TRa-25179", 1312, 13);
  C_Date("1100-1200", 1150, 25);
  Boundary("End Trondheim 12th century");
 };
 R_Date("TRa-25887", 1255, 11);
 R_Date("TRa-25177", 1229, 13);
};
```
