## Supplemental Material 3: Data from calib.org-marine for "Walrus Population-specific Marine Reservoir Offsets (ΔR) for Calibration of Radiocarbon Dates: Implications for Arctic Chronologies and Medieval Trade"

Supplemental Material 3. Predominantly shell-derived ΔR values from areas relevant to the walrus populations considered in this paper, from the 14CHRONO Marine20 Reservoir database at http://calib.org/marine/ (Reimer & Reimer 2001).

### Samples named "Baffin Bay", either as Locality or Subregion:

| MapNo | Lat | Lon | ΔR | σ |
|---|---|---|---|---|
| 2638 | 72.0667 | -59.8333 | -76 | 27 |
| 2641 | 72.0667 | -59.8333 | -56 | 42 |
| 2644 | 72.0667 | -59.8333 | 57 | 31 |
| 2676 | 71.1667 | -58.9333 | -80 | 22 |
| 2677 | 70.0000 | -58.6333 | -1 | 23 |
| 782 | 66.1300 | -65.6700 | 77 | 40 |
| 787 | 66.1500 | -65.7300 | -53 | 40 |
| 788 | 66.1500 | -65.7300 | -83 | 70 |
| 2665 | 64.9333 | -66.3000 | -42 | 23 |
| 790 | 62.7300 | -65.4800 | 47 | 20 |
| 781 | 62.8300 | -66.5800 | -3 | 40 |
| 783 | 62.8300 | -66.5800 | 57 | 40 |
| 785 | 62.6600 | -66.8000 | 47 | 40 |
| 2667 | 71.9500 | -73.9333 | 117 | 28 |
| 2668 | 72.1333 | -74.3333 | 220 | 26 |
| 2685 | 72.1333 | -74.3333 | 220 | 31 |
| 786 | 72.4500 | -74.8700 | 60 | 40 |

npts: 17
Weighted Mean ΔR= 31
Uncertainty= 97

Samples named “West Greenland” or “Greenland coastal waters” (which are located on the Western part of Greenland), either as Locality or Subregion (some overlap with Baffin Bay samples):

| MapNo | Lat | Lon | ΔR | σ |
|---|---|---|---|---|
| 2617 | 74.6167 | -62.8500 | -44 | 26 |
| 2616 | 74.8667 | -62.2000 | -74 | 26 |
| 2638 | 72.0667 | -59.8333 | -76 | 27 |
| 2641 | 72.0667 | -59.8333 | -56 | 42 |
| 2644 | 72.0667 | -59.8333 | 57 | 31 |
| 2615 | 75.4333 | -62.4333 | 86 | 25 |
| 2676 | 71.1667 | -58.9333 | -80 | 22 |
| 2677 | 70.0000 | -58.6333 | -1 | 23 |
| 2618 | 73.2000 | -58.1333 | -8 | 27 |
| 2626 | 66.5500 | -61.8333 | 81 | 26 |
| 2649 | 72.5333 | -57.6000 | -18 | 32 |
| 2619 | 68.2833 | -58.2333 | -56 | 25 |
| 2610 | 76.1500 | -68.4667 | -29 | 20 |
| 2648 | 73.9167 | -56.6667 | -92 | 28 |
| 666 | 76.5500 | -68.7700 | 34 | 52 |
| 40 | 76.5700 | -68.8000 | -190 | 57 |
| 39 | 72.7800 | -56.1700 | -189 | 44 |
| 988 | 76.5600 | -68.9200 | -153 | 60 |
| 2665 | 64.9333 | -66.3000 | -42 | 23 |
| 2647 | 76.4167 | -69.6333 | -17 | 24 |
| 2659 | 76.4167 | -69.6667 | -66 | 25 |
| 2681 | 69.6833 | -55.5000 | -136 | 27 |
| 2608 | 66.5833 | -56.6333 | -26 | 24 |
| 2609 | 66.5833 | -56.6333 | -43 | 31 |

| | | | | |
|---|---|---|---|---|
| 2620 | 66.5833 | -56.6333 | -32 | 24 |
| 2639 | 66.5833 | -56.6333 | -72 | 27 |
| 2687 | 69.6833 | -55.0167 | -131 | 32 |
| 38 | 72.3700 | -54.7300 | -207 | 44 |
| 2630 | 71.3500 | -54.4833 | -72 | 29 |
| 2634 | 71.3500 | -54.4833 | -55 | 25 |
| 2650 | 71.3500 | -54.4833 | -13 | 28 |
| 2680 | 71.3500 | -54.4833 | -93 | 27 |
| 986 | 71.3500 | -54.4800 | 56 | 50 |
| 989 | 71.3500 | -54.4800 | -24 | 20 |
| 2636 | 68.0000 | -54.8333 | -83 | 23 |
| 2614 | 77.0833 | -71.2167 | -52 | 24 |
| 2624 | 68.0000 | -54.5000 | -172 | 26 |
| 36 | 69.2500 | -53.5000 | -309 | 71 |
| 2667 | 71.9500 | -73.9333 | 117 | 28 |
| 2672 | 71.9500 | -73.9333 | 224 | 30 |
| 2688 | 69.2833 | -52.8333 | 24 | 27 |
| 2628 | 65.5667 | -54.5167 | -87 | 29 |
| 2629 | 65.5667 | -54.5167 | -109 | 30 |
| 2668 | 72.1333 | -74.3333 | 220 | 26 |
| 2673 | 72.1333 | -74.3333 | 246 | 27 |
| 2685 | 72.1333 | -74.3333 | 220 | 31 |
| 37 | 69.6800 | -52.0000 | -129 | 58 |
| 665 | 69.5000 | -52.0000 | -85 | 51 |
| 987 | 69.0000 | -52.0000 | 116 | 40 |
| 990 | 69.0000 | -52.0000 | -4 | 25 |
| 2690 | 65.1833 | -53.5500 | -103 | 24 |
| 2664 | 65.4167 | -52.8333 | -81 | 27 |

| | | | | |
|---|---|---|---|---|
| 35 | 69.2200 | -51.1300 | -248 | 72 |
| 2627 | 62.1000 | -55.0000 | -149 | 27 |
| 2625 | 63.5833 | -52.8500 | -100 | 21 |
| 2651 | 64.0333 | -52.4167 | 13 | 28 |

npts: 56
Weighted Mean ΔR= -30
Uncertainty= 97

### Samples named “Foxe basin”, either as Locality or Subregion

| MapNo | Lat | Lon | ΔR | σ |
|---|---|---|---|---|
| 816 | 63.6800 | -80.2000 | -43 | 40 |
| 814 | 62.9500 | -81.8300 | -93 | 40 |
| 796 | 63.0000 | -82.6500 | 87 | 50 |
| 815 | 63.0000 | -82.6500 | -13 | 40 |
| 817 | 63.0000 | -82.6500 | -73 | 50 |
| 824 | 62.9800 | -82.6800 | -3 | 40 |
| 797 | 63.6000 | -82.0000 | -123 | 50 |
| 818 | 63.6000 | -82.0000 | 67 | 50 |
| 792 | 64.4000 | -77.9300 | 157 | 50 |
| 826 | 64.4000 | -77.9300 | 87 | 50 |
| 821 | 66.9200 | -81.3300 | 207 | 40 |
| 2041 | 67.8622 | -77.2033 | 82 | |
| 2042 | 67.8622 | -77.2033 | 52 | |
| 2043 | 67.8622 | -77.2033 | -8 | |
| 819 | 69.2700 | -77.8000 | 97 | 40 |
| 794 | 66.4700 | -86.2000 | 87 | 50 |
| 832 | 66.5200 | -86.2500 | 162 | 20 |
| 804 | 69.4000 | -80.8800 | 107 | 50 |
| 807 | 69.3100 | -81.5900 | 377 | 50 |

| | | | | |
|---|---|---|---|---|
| 823 | 69.3100 | -81.5900 | 117 | 50 |
| 808 | 69.5700 | -80.2800 | 377 | 50 |
| 798 | 69.3400 | -81.7200 | 177 | 40 |
| 799 | 69.3400 | -81.7200 | 87 | 50 |
| 801 | 69.3400 | -81.7200 | 197 | 40 |
| 803 | 69.3400 | -81.7200 | 127 | 50 |
| 825 | 69.3400 | -81.7200 | 137 | 40 |
| 828 | 69.3400 | -81.7200 | 307 | 50 |
| 829 | 69.3400 | -81.7200 | 127 | 50 |
| 833 | 69.3400 | -81.7200 | 277 | 20 |
| 835 | 69.3400 | -81.7200 | 132 | 20 |
| 806 | 69.3800 | -81.7400 | 297 | 40 |
| 810 | 69.3800 | -81.7400 | 357 | 40 |
| 811 | 69.3800 | -81.7400 | 317 | 50 |
| 813 | 69.3800 | -81.7400 | 307 | 50 |
| 830 | 69.3800 | -81.7400 | 357 | 60 |
| 834 | 69.3800 | -81.7400 | 327 | 20 |
| 793 | 69.6200 | -81.1000 | 177 | 50 |
| 795 | 69.6200 | -81.1000 | 187 | 50 |
| 800 | 69.6200 | -81.1000 | 97 | 50 |
| 831 | 69.6200 | -81.1000 | 217 | 60 |
| 827 | 69.6900 | -80.8800 | 137 | 50 |
| 805 | 69.7300 | -77.6300 | 37 | 50 |
| 822 | 69.9400 | -80.3200 | 67 | 40 |
| 802 | 70.0100 | -80.2000 | 17 | 40 |
| 809 | 70.0100 | -80.2000 | 167 | 50 |
| 812 | 70.0100 | -80.2000 | 167 | 40 |
| 820 | 70.0100 | -80.2000 | 97 | 40 |

npts: 47
Weighted Mean ΔR= 164
Uncertainty= 122

### Hudson Bay and/or Hudson Strait in Locality and/or in Subregion

| MapNo | Lat | Lon | ΔR | σ |
|---|---|---|---|---|
| 872 | 56.4500 | -78.8900 | -183 | 40 |
| 2045 | 63.7255 | -80.2117 | 12 | |
| 871 | 56.5000 | -76.6000 | 26 | 40 |
| 2046 | 63.7525 | -80.1682 | 12 | |
| 2047 | 63.7525 | -80.1682 | -18 | |
| 2048 | 63.7525 | -80.1682 | -8 | |
| 2049 | 63.7525 | -80.1682 | -33 | |
| 2050 | 63.7525 | -80.1682 | -18 | |
| 2051 | 63.7525 | -80.1682 | -28 | |
| 2052 | 63.7525 | -80.1682 | 22 | |
| 2053 | 63.7525 | -80.1682 | -18 | |
| 2054 | 63.7525 | -80.1682 | 17 | |
| 2055 | 63.7525 | -80.1682 | -13 | |
| 2056 | 63.7525 | -80.1682 | -23 | |
| 2060 | 63.6760 | -77.3712 | 56 | |
| 2061 | 63.6760 | -77.3712 | 86 | |
| 2063 | 63.6760 | -77.3712 | 66 | |
| 2064 | 63.6760 | -77.3712 | 91 | |
| 2057 | 62.9882 | -82.2433 | -13 | |
| 2059 | 62.9882 | -82.2433 | -9 | |
| 874 | 56.2500 | -76.3300 | -24 | 50 |
| 875 | 55.2800 | -77.7500 | -33 | 50 |
| 570 | 64.4000 | -77.9300 | 87 | 50 |

| | | | | |
|---|---|---|---|---|
| 879 | 64.2300 | -76.5500 | -153 | 40 |
| 880 | 64.2300 | -76.5500 | -23 | 40 |
| 881 | 64.2300 | -76.5500 | -63 | 50 |
| 2038 | 64.2315 | -76.5422 | 21 | |
| 2039 | 64.2315 | -76.5422 | 101 | |
| 2040 | 64.2315 | -76.5422 | 32 | |
| 876 | 64.3300 | -75.5700 | -3 | 50 |
| 2044 | 61.5972 | -71.9620 | 42 | |
| 2058 | 61.5972 | -71.9620 | 37 | |
| 877 | 61.6300 | -71.9700 | -104 | 50 |
| 2065 | 62.5512 | -70.5950 | 2 | |
| 873 | 58.8300 | -94.0700 | -103 | 40 |

npts: 35
Weighted Mean ΔR= -59
Uncertainty= 78

### Eastern Greenland

| MapNo | Lat | Lon | ΔR | Σ |
|---|---|---|---|---|
| 2622 | 74.5833 | -18.3833 | 7 | 28 |
| 2635 | 74.5833 | -18.3833 | 30 | 26 |
| 2663 | 74.5833 | -18.3833 | 27 | 26 |
| 2683 | 74.5833 | -18.3833 | 98 | 36 |
| 2611 | 74.5000 | -18.6667 | 58 | 26 |
| 20 | 74.3800 | -19.2700 | -23 | 38 |
| 2658 | 75.9167 | -19.3333 | -99 | 22 |
| 2643 | 74.1667 | -20.3333 | 49 | 38 |
| 31 | 76.7500 | -18.7300 | -39 | 58 |
| 2662 | 73.4333 | -21.2167 | -39 | 24 |
| 2675 | 72.7000 | -14.8167 | -21 | 20 |

| | | | | |
|---|---|---|---|---|
| 21 | 73.4700 | -21.5000 | 37 | 47 |
| 22 | 73.4700 | -21.5000 | 7 | 54 |
| 2660 | 70.4500 | -20.3667 | -77 | 26 |
| 2693 | 72.7500 | -22.9333 | 115 | 24 |
| 2656 | 73.2667 | -23.2500 | 131 | 27 |
| 2691 | 73.2667 | -23.2500 | 163 | 23 |
| 2692 | 73.2667 | -23.2500 | 1141 | 27 |
| 27 | 70.8200 | -22.4800 | -149 | 52 |
| 2653 | 70.8333 | -22.5167 | 173 | 25 |
| 2655 | 70.8333 | -22.5167 | 240 | 38 |
| 2682 | 70.8333 | -22.5167 | -25 | 33 |
| 2695 | 70.8333 | -22.5167 | 395 | 29 |
| 26 | 70.4500 | -22.2300 | -2 | 74 |
| 23 | 70.8300 | -22.5500 | 27 | 39 |
| 2694 | 70.7167 | -22.4833 | 164 | 23 |
| 2698 | 70.7167 | -22.4833 | 64 | 38 |
| 791 | 70.5000 | -22.5000 | -194 | 60 |
| 2654 | 70.4500 | -22.5833 | 18 | 26 |
| 2621 | 66.8333 | -20.0333 | -38 | 25 |
| 25 | 69.4200 | -24.0800 | -85 | 73 |
| 2696 | 73.2500 | -25.7000 | 716 | 30 |
| 2642 | 72.7167 | -26.6333 | -43 | 27 |
| 2633 | 73.1000 | -27.2833 | -22 | 35 |
| 2652 | 73.1000 | -27.2833 | -29 | 31 |
| 2671 | 69.2167 | -8.3833 | -17 | 24 |
| 2670 | 66.3833 | -7.4167 | -49 | 28 |
| 2686 | 65.6667 | -35.5333 | -52 | 24 |

npts: 38
Weighted Mean ΔR= 84

Uncertainty= 244

### Iceland

| MapNo | Lat | Lon | ΔR | Σ |
|---|---|---|---|---|
| 1847 | 66.0000 | -18.0000 | -23 | 45 |
| 1848 | 66.0000 | -18.0000 | -44 | 23 |
| 65 | 66.0000 | -17.5000 | -164 | 34 |
| 1982 | 66.5265 | -18.1957 | -156 | 43 |
| 67 | 66.2500 | -15.3200 | -53 | 59 |
| 60 | 65.1700 | -22.0000 | -108 | 35 |
| 66 | 64.3300 | -22.0000 | -334 | 58 |
| 57 | 64.0000 | -22.0000 | -60 | 51 |
| 58 | 64.0000 | -22.0000 | 18 | 51 |
| 59 | 64.0000 | -22.0000 | 57 | 51 |
| 61 | 64.6700 | -14.2500 | -135 | 41 |
| 64 | 64.1200 | -22.2000 | -103 | 42 |
| 62 | 65.2800 | -14.0000 | -124 | 41 |
| 63 | 64.3300 | -22.5000 | -134 | 35 |

npts: 14
Weighted Mean ΔR= -94
Uncertainty= 76

### Samples from the archipelago of Franz Josef Land

| MapNo | Lat | Lon | ΔR | σ |
|---|---|---|---|---|
| 615 | 81.2300 | 53.8500 | -283 | 40 |
| 616 | 79.9200 | 49.8000 | -265 | 48 |

npts: 2
Weighted Mean ΔR= -276
Uncertainty= 31

### Pacific: samples that are in the general area around the coordinates of our two Pacific walrus

| MapNo | Lat | Lon | ΔR | σ |
|---|---|---|---|---|
| 705 | 70.4000 | -161.4200 | 454 | 40 |
| 702 | 70.4000 | -161.4200 | 154 | 50 |
| 715 | 70.4000 | -161.4200 | 164 | 25 |
| 945 | 65.2500 | -166.6700 | 194 | 50 |
| 959 | 65.2500 | -166.6700 | 369 | 20 |
| 954 | 65.2700 | -166.3700 | 424 | 40 |
| 960 | 65.2700 | -166.3700 | 294 | 20 |
| 706 | 71.4000 | -156.4800 | 264 | 40 |
| 707 | 71.4000 | -156.4800 | 404 | 30 |
| 704 | 71.4000 | -156.4800 | 314 | 40 |
| 712 | 71.4000 | -156.4800 | 404 | 60 |
| 147 | 55.5000 | -162.0000 | 97 | 50 |

npts: 12
Weighted Mean ΔR= 305
Uncertainty= 102
